# Correlated protein-RNA associations and a requirement for HNRNPU in long-range Polycomb recruitment by the lncRNAs *Airn* and *Kcnq1ot1*

**DOI:** 10.1101/2025.07.16.665154

**Authors:** McKenzie M. Murvin, Shuang Li, Elizabeth W. Abrash, Bridget A. Peck, Samuel P. Boyson, Zhiyue Zhang, Rachel E. Cherney, J. Mauro Calabrese

## Abstract

The lncRNAs *Airn* and *Kcnq1ot1* recruit Polycomb Repressive Complexes (PRCs) and repress genes over multi-megabase genomic intervals, but how they interact with proteins to direct repression remains poorly understood. We conducted formaldehyde-linked RNA-immunoprecipitations of 27 proteins from trophoblast stem cells, using a protocol exhibiting similar signal-to-noise and post-lysis reassociation ratios as CLIP and CLAP. Patterns of protein associations across *Airn* and *Kcnq1ot1* were more similar to each other than to nearly all other transcripts and partitioned to extents that mirrored degree of repression each lncRNA induced, implying connections to mechanism. Indeed, HNRNPU, an essential protein that helps *Xist* localize to chromatin, was enriched within and required to direct PRC-catalyzed modifications by *Airn* and *Kcnq1ot1,* without being required for their localization. Our study provides insights into the role of HNRNPU in lncRNA-mediated gene regulation and reports architectures of protein association across *Airn* and *Kcnq1ot1* relative to the transcriptome at large.

**Author summary:** Mammalian genomes express thousands of long noncoding RNAs (lncRNAs). While most remain functionally uncharacterized, a handful are known to regulate genes via epigenetic pathways. Whether or not these known regulatory lncRNAs operate through shared or divergent mechanisms remains unclear. To gain insight, we compared RNA-protein associations across three known regulatory lncRNAs -- *Airn*, *Kcnq1ot1*, and *Xist* -- relative to the broader transcriptome, using a protocol to immunoprecipitate RNA from formaldehyde-crosslinked cells. Benchmarking of our RNA-immunoprecipitation protocol against methods called CLIP and CLAP revealed similar results and complementary strengths among methods. We discovered similarities in RNA-protein associations between *Airn* and *Kcnq1ot1* that nominated them as mechanistically important. One such protein, HNRNPU, was required for *Airn* and *Kcnq1ot1* to coordinate long-range recruitment of Polycomb complexes, without giving the appearance of being required to tether either lncRNA to chromatin, as HNRNPU has been proposed to do for *Xist*. Our study identifies RNA-protein associations that correlate with long-range gene regulation by lncRNAs and provides new perspectives on the potential mechanisms through which HNRNPU participates in lncRNA-mediated epigenetic control. We also benchmark a protocol to recover RNA-protein interactions that is simple to execute and has complementary strengths relative to other methods.

## Introduction

Many RNAs produced by mammalian genomes are long and have little potential to encode protein even after splicing (long noncoding RNAs; lncRNAs). While most lncRNAs are functionally unannotated, several play important roles in health by regulating gene expression *in cis*, on the same allele from which they are transcribed.^1^ The sequence-based logic that lncRNAs employ to carry out regulatory functions remains poorly understood, leaving much unclear about mechanism and the extent to which uncharacterized lncRNAs regulate gene expression.

The best-studied and most potent *cis*-acting regulatory lncRNA is *Xist*, which silences gene expression over the 165 megabase (Mb) X-chromosome.^2^ On autosomes, at least two lncRNAs also repress gene expression *in cis*: *Airn* and *Kcnq1ot1*. Both are imprinted and expressed from paternally inherited alleles, where they can repress genomic regions spanning ∼15 Mb and ∼3 Mb around their sites of transcription, respectively.^3–8^ While *Xist* is 18 kilobases (kb) long, robustly spliced, and abundant, *Airn* and *Kcnq1ot1* are upwards of 90 kb long, predominantly unspliced, and lower in abundance.^4,9–14^ In female mouse trophoblast stem cells (TSCs), which express all three lncRNAs, we have observed that at the steady state, *Xist* accumulates to about 230 copies per cell, whereas *Airn* and *Kcnq1ot1* each accumulate to only 7 or 8 copies, a 25 to 30-fold difference.^4^

Despite their differences, *Airn*, *Kcnq1ot1,* and *Xist* require several of the same proteins to carry out repression. These include the histone-modifying enzymes Polycomb Repressive Complex 1 (PRC1), which catalyzes monoubiquitination of lysine 119 on Histone H2A; Polycomb Repressive Complex 2 (PRC2), which catalyzes trimethylation of lysine 27 on Histone H3; and G9a/EHMT2, which catalyzes the demethylation of lysine 9 on Histone H3.^15–23^ Additionally, all three lncRNAs require the RNA-binding protein (RBP) HNRNPK to recruit PRCs, and *Xist* and *Kcnq1ot1* appear to require the RBP SPEN to induce gene silencing.^4,13,24^

It remains unclear whether *Airn*, *Kcnq1ot1,* and *Xist* engage with additional shared cofactors or different ones to maintain repression. It also remains unclear whether the lncRNAs exhibit distinct properties in relation to the broader transcriptome or even to each other. Although protein interactions within *Xist* have been characterized,^2,25–38^ the RBPs that interact with *Airn* and *Kcnq1ot1*, their patterns of interaction within each lncRNA, and the functional relevance of these interactions are incompletely defined. In addition to providing insights into mechanism, addressing these unknowns could help identify predictive features of mammalian regulatory RNAs and develop a better understanding of the functions of lncRNA-associated RBPs, many of which are essential for health, such as HNRNPU.^39–43^

Herein, we used a formaldehyde-based RNA immunoprecipitation-sequencing (RIP-seq) protocol^44^, which we demonstrate exhibits similar signal-to-noise and post-lysis RNA association ratios as crosslinking immunoprecipitation (CLIP) and crosslinking affinity purification (CLAP;^35^), to examine protein associations across *Airn*, *Kcnq1ot1,* and *Xist* in TSCs. We found that protein associations within *Airn* and *Kcnq1ot1* were more correlated to each other than to *Xist* and essentially all other chromatin-enriched RNAs, and that the extent to which the associations partitioned into communities correlated with the degree of repression induced by each lncRNA. HNRNPU, an RBP necessary for *Xist*’s localization to the X chromosome, exhibited enriched associations with *Airn* and *Kcnq1ot1* and was required to maintain their Polycomb-repressed domains, yet was not required for their proper localization. Our study highlights a RIP protocol that offers advantages and complementary insights relative to CLIP and CLAP, provides a roadmap for interrogation of RNA-protein associations across *Airn*, *Kcnq1ot1*, and other chromatin-associated RNAs, and offers new insights into gene regulation by HNRNPU.

## Results

### A formaldehyde-based RIP protocol returns similar signal-to-noise and post-lysis reassociation ratios as CLIP and CLAP

*Xist* directly or indirectly associates with hundreds of proteins, many of which bind RNA, are expressed at near-micromolar concentrations, and regulate many processes, including splicing, RNA export, and translation.^25–34,45,46^ Prior analysis by formaldehyde-based RIP revealed that one such protein, HNRNPK, exhibited enriched associations not only within *Xist* but also within *Airn* and *Kcnq1ot1,* in patterns implicating HNRNPK and its RNA-bound regions in PRC recruitment.^4,13^ We reasoned that other proteins important for biogenesis or repressive function of *Airn*, *Kcnq1ot1*, or *Xist* might also exhibit distinct patterns of enrichment within the lncRNAs.

To investigate, we performed RIP-seq from formaldehyde-crosslinked, female, F1-hybrid mouse TSCs examining a targeted panel of 25 different RBPs, two components of PRC1 (RING1B and RYBP), and six biological replicates of non-specific IgG control (Figure 1A; Table S1;^25–34^). We focused our panel of RIPs on proteins previously found to be associated with *Xist*, because like *Xist*, *Airn* and *Kcnq1ot1* are also chromatin-associated and might draw from a similar pool of proteins to enact repression. *Airn* and *Kcnq1ot1* are also ∼25-fold lower in abundance than *Xist*,^4^ rendering proteomics-based approaches to identify RNA-protein interactomes more challenging to execute and less suitable for direct comparisons.^47^ We performed experiments in F1-hybrid TSCs, where SNPs between parental alleles enabled us to monitor effects of each lncRNA *in cis*.^4,14,48^ Our approach enabled mapping of patterns of protein association in *Airn*, *Kcnq1ot1*, and *Xist* relative to the transcriptome at large, including direct and protein-bridged associations that may underlie higher-order assemblies.^13,44,49–51^ Because formaldehyde captures direct and indirect (protein-bridged) RNA-protein interactions, we use the term “association” rather than “binding” to describe RNA-protein interactions detected. Like other approaches to detect RNA-protein interactions, formaldehyde-based RIP recovers non-specific associations, which we model below using IgG and non-specific antibody controls.

**Figure 1.**
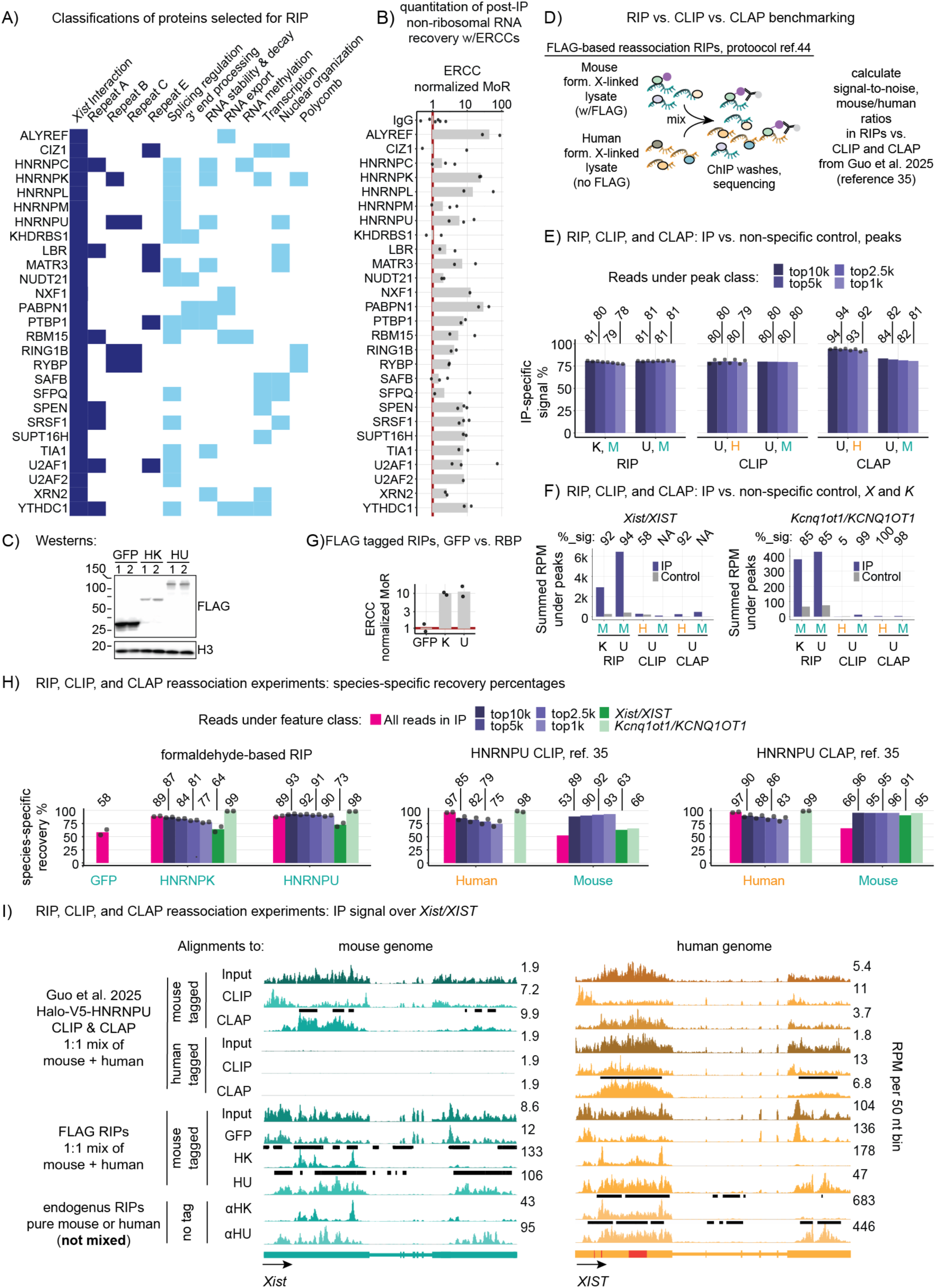
A formaldehyde-based RIP protocol returns similar signal-to-noise and post-lysis reassociation ratios as CLIP and CLAP. **(A)** 27 proteins selected for RIP-seq in TSCs, along with classifications. “*Xist* Interaction,” *Xist*-associated protein; “Repeat A/B/C/E,” protein enriched over *Xist*-Repeat.^25–34^ **(B)** Normalized MoR of RIP replicates relative to average IgG. **(C)** FLAG westerns. (C-I): HK/K, HNRNPK; HU/U, HNRNPU. **(D)** RIP reassociation test overview. **(E)** IP-specific signal percentages by peak class. **(F)** RPM under peaks within *Xist/XIST, Kcnq1ot1/KCNQ1OT1*. %_sig, 100*IP/(IP + control). (**G**) Normalized MoR for FLAG-RBP RIPs relative to FLAG-GFP. (**H**) Species-specific recovery percentages (percent of reads or RPKM aligning to tag-expressing vs. non-expressing genome). (**I**) Wiggle density profiles of RIP, CLIP, and CLAP data over *Xist/XIST*. Black bars, peaks called in sample. Red bars, Repeats B/D. See also Figures S1-4, Table S1.

RIPs for all but two of 27 proteins (NXF1 and U2AF2) were performed in at least biological duplicate (Table S1). To identify RNA regions enriched over IgG control, we employed a peak calling strategy that requires peaks to exhibit a >2-fold increase in signal in ≥2 replicates over averaged IgG control (for the 25 RIPs replicated in this study). This threshold was empirically selected to reduce false positives due to noise and false negatives due to high background over certain regions, such as those associated with the splicing machinery.^13,44,52–54^ Read densities suggest the minimum resolution of our protocol to be 200-300 nucleotides; we set our minimum peak size to be 200 nucleotides, accordingly.^44^ As expected, read densities under peaks were highly correlated between replicates (Table S1). Likewise, in 15 of 19 cases where motifs were previously defined, sequences under peaks were enriched for expected motifs (Figures S1). In remaining cases, antibody specificity was confirmed by western blot (Figure S2). We observed expected patterns of enrichment for several RBPs over known binding regions in *Xist*, additional evidence that the protocol is recovering expected RNA-protein associations (Figure S3;^2,35–38^).

We next sought to estimate amount of RNA enriched by each RIP relative to IgG control. As part of our standard protocol, we add ERCC Spike-In control RNAs to immunoprecipitated RNA prior to library preparation. After sequencing, ERCC-counts can be used to infer amount of non-ribosomal RNA recovered by RIP; the lower the ERCC-counts, the greater the fraction of non-ribosomal RNA recovered.^44,55^ Across all RIPs, we subjected ERCC-counts to median of ratios normalization and inverted values to facilitate comparisons to IgG (“normalized MoR”; higher values, more RNA). 54 out of 59 individual RIPs returned MoR values greater than that of averaged IgG control (Figure 1B; Table S1). Even between replicates of the same RIP that recovered more and less RNA than IgG control, respectively (e.g. Ciz1), signal patterns were still highly correlated (Pearson’s r., >0.86; Table S1). These analyses underscore the general success of our protocol and document which RIPs returned the highest levels of overall signal relative to IgG control.

Recent work has promoted CLAP as an alternative to RIP and CLIP, premising that CLAP employs covalent-epitope-tagging and denaturing washes to reduce background in post-lysis protein-recovery steps.^35^ It has also been suggested that at least one native RIP protocol can be confounded by post-lysis RNA-protein reassociations.^56^ Therefore, we sought to compare signal-to-noise and post-lysis RNA-protein association in our version of RIP^44^ versus CLIP and CLAP^35^. We stably introduced doxycycline-inducible 3xFLAG-tagged cDNAs of nuclear-localized GFP, HNRNPK, and HNRNPU – the latter being *Xist*-associated RBPs studied in more depth below – into mouse embryonic stem cells (ESCs) that also express *Xist.*^53,57^ We induced expression of *Xist* and FLAG-tagged proteins, formaldehyde-crosslinked the ESCs and 293T human cells, mixed them in a 1:1 ratio, then performed FLAG RIP (Figures 1C,D). We also performed HNRNPK and HNRNPU RIP from unmixed 293T extracts using non-FLAG antibodies (same antibodies used below), to identify 293T peaks (Figure S4). We examined signal-to-noise and species-specific RNA recovery ratios in RIPs versus those from Halo-V5-tagged HNRNPU CLIP and CLAP performed in reciprocally mixed *Xist*-expressing mouse SM33-ESCs and human 293Ts (Figure S4;^35^). Data analyzed can be viewed in Table S1 and (https://genome.ucsc.edu/s/mmurvin/mm10_12_18_25_RIP_reassoc_CLIP_CLAP_ChIP_RNA; https://genome.ucsc.edu/s/mmurvin/hg38_RIP_reassoc_01_08_25; https://data.mendeley.com/preview/3b3xys2743?a=622e79bc-637f-4c7f-9506-31b022f7a155).

We first analyzed signal over control-inferred noise. For all three methods, peaks were defined as above, requiring 2x greater signal in both replicates relative to non-specific control (when replicates were performed;^35^). FLAG-GFP served as non-specific control for FLAG RIPs and tag-lacking datasets as controls for CLIP and CLAP (Figure S4;^35^). We used CLAP peaks to analyze CLIP data to simplify comparisons and because high variation between human CLIP replicates prevented clear peaks from being called.^35^ We calculated RPM-normalized signal-under peak values for IPs and non-specific controls (Figure S4), then calculated “IP-specific signal” percentages, defined as 100 * [IP_RPM_under_peak]/([IP_RPM_under_peak]+[control_RPM_under_peak]). We ranked peaks using the product of IP_RPM_under_peak * IP-specific signal. Lastly, we summed RPM values to calculate aggregate IP-specific signal percentage for different classes of peaks. Across classes, IP-specific signal percentages were highest in human CLAP (92-94) and lower but similar between RIP, CLIP, and mouse CLAP (RIP/CLIP, 78-81; mouse CLAP, 81-84; Figure 1E).

We next examined signal and control-inferred noise within *Xist/XIST* and *Kcnq1ot1/KCNQ1OT1* because of their relevance to our study. *Airn* is neither robustly expressed in ESCs nor conserved in humans, precluding its analysis here. We summed IP_RPM_under_peak and control_RPM_under_peak values over the length of *Xist/XIST* and *Kcnq1ot1/KCNQ1OT1* and examined IP-specific signal. We dropped *Xist* from CLIP and CLAP analyses because it was not expressed in the mouse mixed with human-tagged samples that serve as non-specific controls for mouse-tagged samples (Figures 1I, S4;^35^). Across methods, IP_RPM_under_peak values were ∼10-100-fold higher in RIP versus CLIP and CLAP, perhaps reflecting increased crosslinking efficiency of formaldehyde relative to UV light (Figure 1F). Comparing our lncRNAs of interest, IP-specific signal percentages in RIP ranged between 85 and 94, covered a broad range in CLIP, and ranged from 92 to 100 in CLAP (Figure 1F). Normalized MoR values were 10-fold higher in FLAG-HNRNPK and FLAG-HNRNPU versus FLAG-GFP RIPs (Figure 1G), implying the RBP-RIPs recovered 10-times the amount of non-ribosomal RNA as non-specific control, and that their absolute IP-specific signal percentages may be higher than those calculated using the proportion-based RPM.

We next examined post-lysis reassociation in RIP, CLIP, and CLAP by considering read alignments within combined mouse-and-human genomes. We specify “species-specific recovery” to describe the percentage of reads uniquely aligning to the genome expressing the epitope-tagged protein over the sum of reads uniquely aligning to both genomes (epitope-tag-expressing plus not-expressing). Comparing all reads sequenced in each method (including those not within peaks), species-specific recovery percentages in FLAG-HNRNPK and FLAG-HNRNPU RIPs were 89, falling toward the upper range detected by HNRNPU CLIP and CLAP (53-97; Figure 1H). Across the same peak classes from Figure 1E, species-specific recovery percentages remained relatively constant in mouse CLIP and CLAP but gradually decreased in RIPs and human CLIP and CLAP (Figure 1H). The decrease was particularly apparent in RIP peaks within *Xist/XIST*, where FLAG-tagged HNRNPK expressed in mouse clearly recovered known HNRNPK-binding regions in human *XIST* (Repeats B, D^58^; *Xist* species-specific percentage, 64; Figures 1H,I). In contrast, peaks within the more-lowly expressed *Kcnq1ot1/KCNQ1OT1* exhibited species-specific recovery percentages of 95 or greater across all datasets except mouse CLIP (Figure 1H, S4D).

Thus, our formaldehyde-based RIP protocol recovers expected motifs across the transcriptome, known RNA-protein interactions in *Xist*, more non-ribosomal RNA than IgG control, and by our analyses, signal-to-noise and post-lysis reassociation ratios that are comparable with CLIP and CLAP. CLAP generally although not always recovers less noise, whereas RIP can recover higher signal intensity under peak regions, presumably due to efficiency of formaldehyde crosslinking. Across all methods, different RNA targets experienced varying levels of post-lysis reassociation, in a manner we presume scales with affinity and abundance of the RNA targets and cognate RBPs.

### Protein association profiles across *Airn* and *Kcnq1ot1* closely resemble each other and exhibit similarity to intron-containing transcripts from lncRNA and protein-coding genes

Having benchmarked our formaldehyde-linked RIP protocol,^44^ we next sought to determine the extent to which our selected proteins associated with *Airn*, *Kcnq1ot1*, and *Xist* above control and relative to other chromatin-enriched transcripts. To determine significance of enrichment for each protein across each chromatin-enriched transcript, we summed background-corrected signal under peaks for individual replicates of each protein over the length of each chromatin-enriched transcript, then used Wilcoxon rank testing to compare signal for replicate RIPs to six replicates of non-specific IgG control (Table S2). We also ranked the set of chromatin-enriched transcripts by overall levels of association with each protein, enabling transcriptome-wide comparisons (Figure 2A;^13^). Significant associations are denoted as purple-shaded boxes with black text in Figure 2A, and individual replicate values for select proteins and IgG control are shown in 2B. RIP signal was significantly higher than IgG control for 23, 24, and 16 proteins over *Airn*, *Kcnq1ot1*, and *Xist*, respectively (Figure 2A; Table S2). Relative to their own expression levels (assessed by Total and Chromatin-associated RNA-seq), *Airn* and *Kcnq1ot1* exhibited enriched associations with many of the same RBPs, some of which were also enriched within *Xist* (e.g. HNRNPU, RBM15, YTHDC1; Figure 2A). Others exhibited enrichments specific to *Airn* and *Kcnq1ot1* (HNRNPL, SAFB, XRN2; Figure 2A).

**Figure 2.**
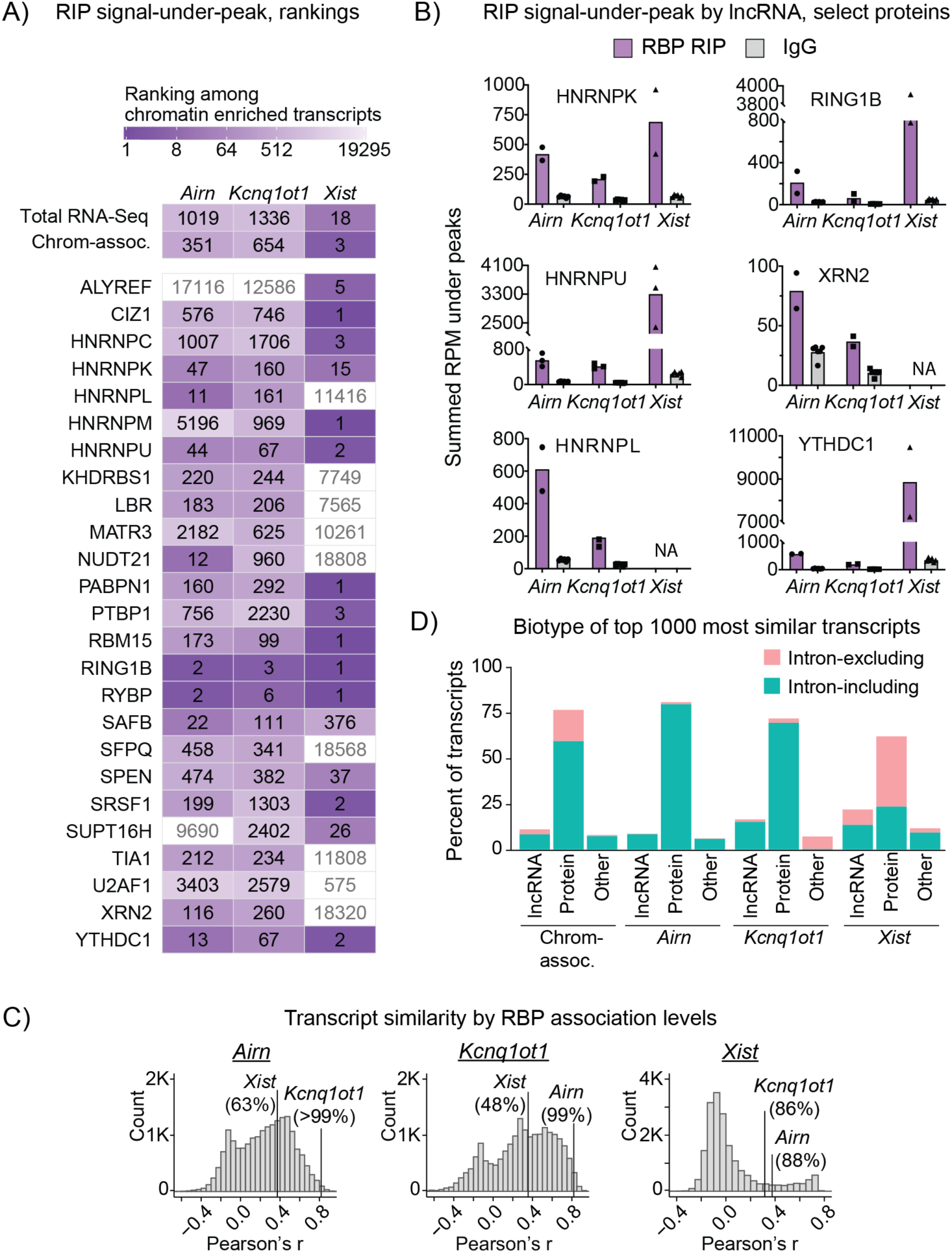
Protein association profiles across *Airn* and *Kcnq1ot1* closely resemble each other and exhibit similarity to intron-containing transcripts from lncRNA and protein-coding genes. (**A**) Signal-under-peak ranks within *Airn*, *Kcnq1ot1*, and *Xist*. “Total RNA-seq”, “Chrom-assoc.”: rankings from total RNA-seq, chromatin-associated RNA-seq.^13^ Black/grey numbers, significant/non-significant associations (p<0.05 Wilcoxon rank). U2AF2, NXF1 RIPs were excluded from significance analyses (singly replicated). (**B**) RIPs versus IgG, signal-under-peaks by lncRNA. “NA”, no peak in lncRNA. **(C)** Pearson’s r histograms comparing signal-under-peak in lncRNAs vs chromatin-enriched transcripts. Labels, percentile of similarity ranking. **(D)** GENCODE genic biotype and intron status of 1000-transcripts most similar to *Airn*, *Kcnq1ot1*, *Xist*.^93^ See also Figures S1-3, Tables S1-3.

We next sought to quantify how similar levels of protein associations across *Airn*, *Kcnq1ot1*, and *Xist* were relative to other chromatin-enriched transcripts. For each transcript, we defined its “protein association profile” as a 27-value vector comprising the summed, background-corrected RIP-seq signal under peaks for that transcript across each protein profiled (Table S2). We then compared association profiles between transcripts using Pearson’s correlation. Of the 19295 chromatin-enriched transcripts in TSCs, 18820 had non-zero background-corrected values for at least one protein examined and were included in comparisons below. Examining these transcripts, association profiles of *Kcnq1ot1* and *Xist* were the 92^nd^ and 6888^th^ most similar to *Airn* (>99^th^ and 63^rd^ percentiles, respectively); association profiles of *Airn* and *Xist* were the 270^th^ and 9744^th^ most similar to *Kcnq1ot1* (99^th^ and 48^th^ percentiles, respectively); and association profiles of *Airn* and *Kcnq1ot1* were the 2352^nd^ and 2601^st^ most similar to *Xist* (88^th^ and 86^th^ percentiles, respectively; Figure 2C; Table S3).

*Airn*, *Kcnq1ot1,* and *Xist* are among the few well-documented *cis*-acting repressive lncRNAs, yet it remains unclear whether they exhibit preferential similarities to protein-coding versus lncRNA genes. To gain insight, we used Pearson’s correlation to identify the top 1,000 transcripts whose protein association profiles were most similar to each lncRNA. In these sets, we evaluated proportions of transcripts produced from lncRNA versus protein-coding genes and compared to proportions from all chromatin-associated RNAs. We also compared proportions of intron-excluding (presumably spliced) and intron-including (presumably nascent) transcripts. These analyses revealed that in terms of protein association profiles, transcripts similar to *Airn*, *Kcnq1ot1,* and *Xist* are mostly produced from protein-coding genes (Figure 2D; >70% in each set). Whereas *Kcnq1ot1*- and *Xist*-similar transcripts exhibited a mild but significant enrichment originating from lncRNA genes, *Airn*-similar transcripts exhibited a mild but significant enrichment originating from protein-coding genes (Figure 2D; p-adj.<0.01; Fisher’s exact). *Airn-* and *Kcnq1ot1*-similar transcripts exhibited significant enrichments for intron-including RNAs, whereas *Xist*-similar transcripts exhibited significant enrichment for intron-excluding RNAs (Figure 2D; p-adj.<0.01; Fisher’s exact).

Thus, the protein association profiles measured over *Airn* and *Kcnq1ot1* are more similar to each other than to the vast majority of other chromatin-enriched RNAs and exhibit a more moderate degree of similarity to *Xist*. Relative to other expressed RNAs, the protein association profiles of *Airn* and *Kcnq1ot1* most frequently resemble intron-containing RNAs produced from protein-coding genes. *Xist* more frequently resembles intron-excluding transcripts, the majority of which are also produced from protein-coding genes.

### Protein association networks in *Airn*, *Kcnq1ot1*, and *Xist* exhibit above average similarities and a correlation between modularity and repressive potency

Our analyses revealed that *Airn* and *Kcnq1ot1* exhibited above-background associations with the majority of *Xist*-associated proteins profiled. We therefore sought to compare the spatial distribution patterns of protein association detected across *Airn*, *Kcnq1ot1*, and *Xist*. Because RIP-seq performed after formaldehyde crosslinking can capture both direct protein-RNA interactions as well as presumed indirect associations through protein-protein contacts, correlated enrichments of different proteins across transcripts might provide clues about coordinated assemblies or functional partnerships, even if not through direct binding. Indeed, discovery of correlated protein associations in different regions of *Xist* has revealed insights into its mechanism.^13,25,37,49,52,59^

We compared background-corrected RIP-seq read density in 25 nucleotide bins for all possible pairs of proteins in all chromatin-enriched transcripts, using Pearson’s correlation. We included all RIP datasets in these analyses to standardize comparisons across the transcriptome. We then used the correlations to create protein association networks in which edge lengths were inversely proportional to correlation coefficients, and node sizes were directly proportional to the number of connected edges. Within networks, communities represent highly correlated nodes that cluster together by the Leiden algorithm.^60^ To enable visual interpretation, Figures 3A and 3B show networks for *Airn*, *Kcnq1ot1*, and *Xist,* along with IgG-subtracted read densities for select proteins color-coded by community.

**Figure 3.**
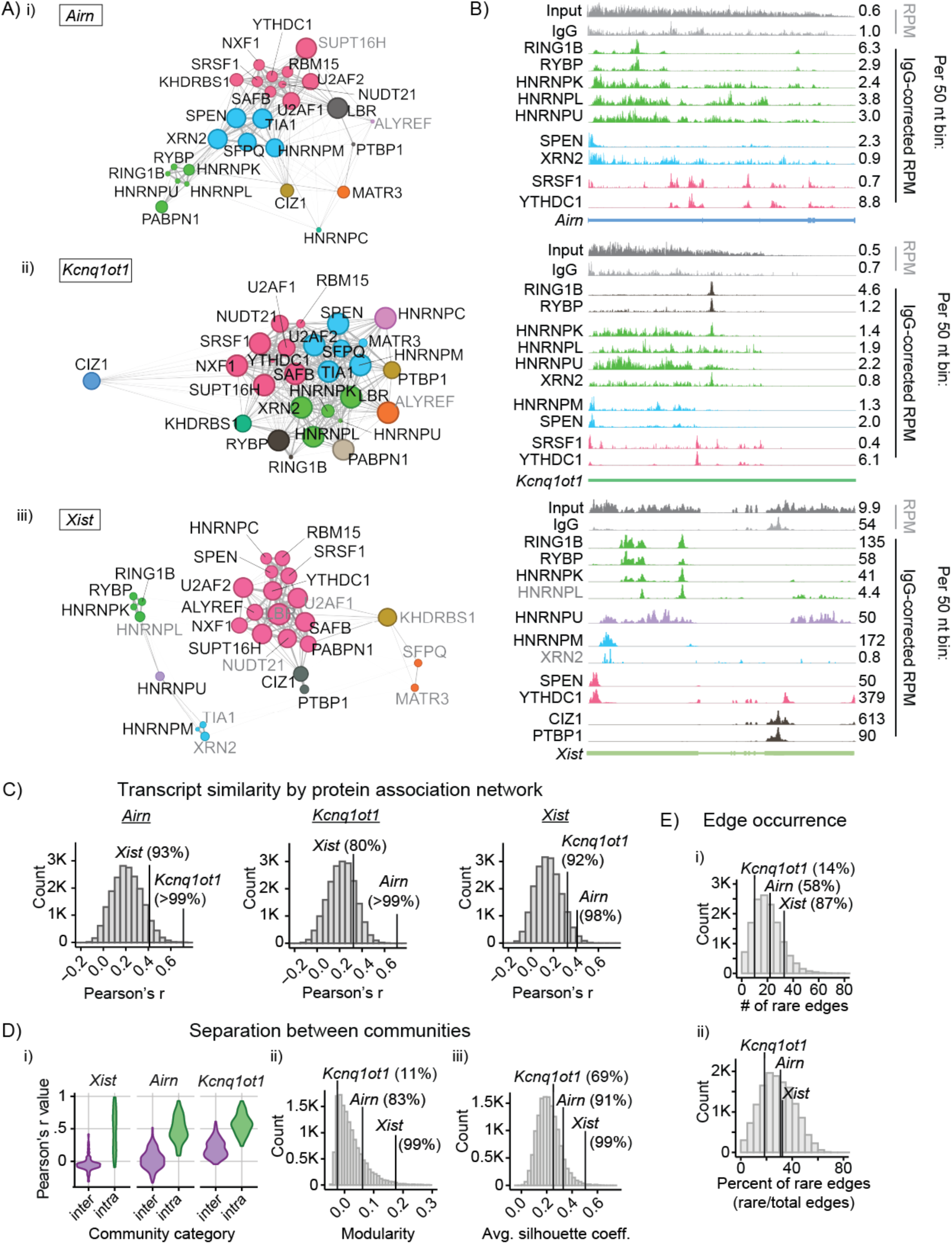
Protein association networks in *Airn*, *Kcnq1ot1*, and *Xist* exhibit above average similarities and a correlation between modularity and repressive potency. (**A**) Network graphs of protein association: (i) *Airn*, (ii) *Kcnq1ot1*, (iii) *Xist*. Edges, nodes connected by r values >0.0. **(B)** Input, IgG, IgG-subtracted RIP-seq density profiles from select communities. Nodes and density profiles: colored by community. Grey text, non-enriched proteins (Figure 2A). **(C)** Pearson’s r histograms comparing association networks in *Airn*, *Kcnq1ot1*, *Xist* relative to other chromatin-enriched transcripts. **(D)** Metrics quantifying separation between communities: (i) Distributions of inter- versus intra-community Pearson’s r; (ii) modularity; (iii) average silhouette distance. **(E)** Number (i) and proportion (ii) of rare edges in *Airn*, *Kcnq1ot1*, *Xist*. See also Tables S3, S4.

We next quantified similarity of protein association networks in *Airn*, *Kcnq1ot1*, and *Xist* relative to all other chromatin-enriched RNAs. Of the 19295 chromatin-enriched RNA transcripts, 19223 had non-zero binned values for at least one protein profiled by RIP-seq and were included below. Relative to 19223 chromatin-enriched transcripts, the association network of *Kcnq1ot1* ranked 6^th^ in similarity to *Airn*, and that of *Airn* ranked 1^st^ in similarity to *Kcnq1ot1*, indicating that patterns of protein association across the two lncRNAs are more similar to each other than to essentially any other chromatin-associated transcript (>99.9^th^ percentiles; Figure 3C; Table S3). The association networks of *Airn* and *Kcnq1ot1* were also more similar to *Xist* than to most other chromatin-associated RNAs, ranking 292^nd^ and 1464^th^, respectively (98^th^ and 92^nd^ percentiles; Figure 3C; Table S3).

We used three orthogonal approaches to examine community structures of protein association networks. First, to quantify the average separation between communities, we examined two distributions within networks: the distribution of Pearson’s r values comparing all proteins within the same communities (intracommunity r values) and the distribution comparing all proteins between different communities (intercommunity r values). While average intracommunity r values were similar between all three lncRNAs, average intercommunity r values were lowest in *Xist*, followed by *Airn*, then *Kcnq1ot1* (Figure 3D, panel (i); p<2e-16, Wilcoxon rank test). This indicates that community structures within *Xist* are more distinct from each other than those within *Airn* and *Kcnq1ot1*. Next, we calculated modularity, a metric to assess how strongly a network partitions into communities relative to a null model. *Xist*’s network had the highest modularity, followed by *Airn*’s, then *Kcnq1ot1*’s, consistent with our intra- vs. inter-community r value analysis (Figure 3D, panel (ii); Table S3). Finally, to assess the degree to which individual nodes (proteins) fit within their assigned community versus other communities, we calculated average silhouette coefficients of all nodes in each transcript’s network.^61^ Higher silhouette coefficients indicate a clearer separation between a node-of-interest’s own community and other communities in the network. Average silhouette coefficients by node were highest in *Xist*, followed by *Airn*, followed by *Kcnq1ot1* (Figure 3D panel (iii); Table S3).

RBPs bind across the transcriptome, and it remains unclear whether correlated associations across *Xist*, *Airn*, and *Kcnq1ot1* occur in other transcripts. To gain insight, we devised a method to quantify which intracommunity connections within *Airn*, *Kcnq1ot1*, and *Xist* represent prevalent versus rare events. Across all chromatin-enriched transcripts, we calculated *p* values describing the likelihood that each possible pair of proteins would be detected in the same community relative to randomized controls. Protein pairs detected more frequently than chance were classified as “prevalent,” and those detected less frequently than by chance, “rare”. Of 351 possible protein pairs, 145 and 191 were classified as prevalent and rare, respectively (p-adj <0.05; Table S4). *Xist* had a higher number of rare intracommunity edges than many other chromatin-associated RNAs (87^th^ percentile), whereas *Airn* and *Kcnq1ot1* had closer to average (58^th^) or lower (14^th^) percentile values, respectively (Figure 3E, panel (i)). About 30% (*Xist* and *Airn*) and 19% (*Kcnq1ot1*) of the edges within the three lncRNAs were classified as rare (Figure 3E, panel (ii)). Notably, edges connecting the PRC1 components RING1B and RYBP to each of the RBPs HNRNPK, HNRNPL, and HNRNPU were rare, whereas edges between any combination of HNRNPK, HNRNPL, and HNRNPU were prevalent, as were edges between PRC1 components and the exoribonuclease XRN2, which was found in communities adjacent to those containing RING1B and RYBP in *Airn* and *Kcnq1ot1* (Table S4).

Thus, the protein association networks within *Airn* and *Kcnq1ot1* were more similar to each other than to essentially all other expressed transcripts, and likewise exhibited similarity to the network within *Xist* that ranked above the 90^th^ percentile. In terms of network structure, communities of association within *Xist* were more separated from each other than those within *Airn*, which were in turn more separated from each other than those within *Kcnq1ot1*. This trend is a striking mirror of the genomic ranges over which each lncRNA represses genes and recruits PRCs; whereas *Xist*’s repressive range spans the entire X chromosome, *Airn*’s spans 15 Mb, and *Kcnq1ot1*’s spans 3 Mb.^4^ Lastly, relative to other chromatin-enriched transcripts, protein association networks of *Airn*, *Kcnq1ot1*, and *Xist* contain both prevalent and rare connected edges, suggesting the lncRNAs engage with proteins in ways that are common in certain instances and uncommon in others. While HNRNPK, HNRNPL, and HNRNPU appear to commonly associate across chromatin-enriched transcripts, their correlated associations with PRC1 are relatively rare, suggesting a connection to repressive mechanism.

### HNRNPU is required for maintenance of PRC-directed modifications induced by *Airn*, *Kcnq1ot1*, and *Xist*

Recognizing Polycomb recruitment as a key function of *Airn* and *Kcnq1ot1*, our attention was drawn to network edges that linked PRC1 components RING1B and RYBP to HNRNPK, HNRNPU, and XRN2, respectively. All three RBPs exhibited enriched associations with *Airn* and *Kcnq1ot1* by RIP-seq (Figure 2B). Whereas HNRNPK is known to be important for PRC recruitment by *Airn* and *Kcnq1ot1,*^4,13,34^ HNRNPU and XRN2 have not been investigated vis-a-vis *Airn* or *Kcnq1ot1*. However, prior data suggest that HNRNPU is required for *Xist* function, presumably stabilizing its association with chromatin,^62–68^ and XRN2 has been proposed to promote H3K27me3 accumulation on autosomes. ^32,69,70^

To determine whether HNRNPU or XRN2 is required for repression by *Airn* and *Kcnq1ot1,* we depleted each protein in TSCs using doxycycline-inducible CRISPR-Cas9 technology.^71^ We additionally depleted YTHDC1, a protein exhibiting enriched association with *Airn*, *Kcnq1ot1,* and *Xist* (Figures 2A,B), which may also interact with PRC2 and promote silencing by *Xist.*^72–74^ As a positive control, we depleted the PRC1-bridging factor HNRNPK.^4,13^ As a negative control, we delivered a non-targeting (NTG) sgRNA along with the same doxycycline-inducible Cas9 construct.^14^ Following four days of Cas9 induction (“(+) dox”), levels of targeted proteins were substantially depleted (Figures 4A,B, S7). To examine whether protein depletion altered PRC1-and PRC2-directed modifications (H2AK119ub and H3K27me3, respectively) in lncRNA-targeted regions, we performed ChIP-seq and analyzed SNP-overlapping reads. These analyses enabled us to distinguish effects on paternally-inherited “B6” alleles, on which the lncRNAs are expressed, versus maternally-inherited “CAST” alleles, on which the lncRNAs are silent.^4,14,48^ As expected, in cells expressing NTG sgRNA, H2AK119ub and H3K27me3 were unchanged or, in the case of H2AK119ub over the inactive X-chromosome, changed significantly but modestly, by ∼10% (Figure S5). Also as expected, in cells depleted of HNRNPK, H2AK119ub and H3K27me3 were significantly reduced in the *Airn*, *Kcnq1ot1,* and *Xist* target regions (Figures 4C, S6, S7). Depletion of XRN2 or YTHDC1 did not reduce H2AK119ub or H3K27me3 in the *Airn* or *Kcnq1ot1* target regions (Figure S8). Over the inactive X-chromosome, depletion of XRN2 led to minimal but statistically significant changes, and depletion of YTHDC1 led to significant but more moderate changes than those observed after depletion of HNRNPK (Figures S7, S8). In contrast, depletion of HNRNPU clearly and significantly reduced H2AK119ub and H3K27me3 in the *Airn*, *Kcnq1ot1*, and *Xist* target regions, and in the *Airn* target domain, the changes exceeded those observed upon depletion of HNRNPK (Figures 4D, S7). We conclude that HNRNPU is required to maintain PRC-directed histone modifications not only over the X-chromosome,^65^ but also in genomic regions repressed by *Airn* and *Kcnq1ot1*.

**Figure 4.**
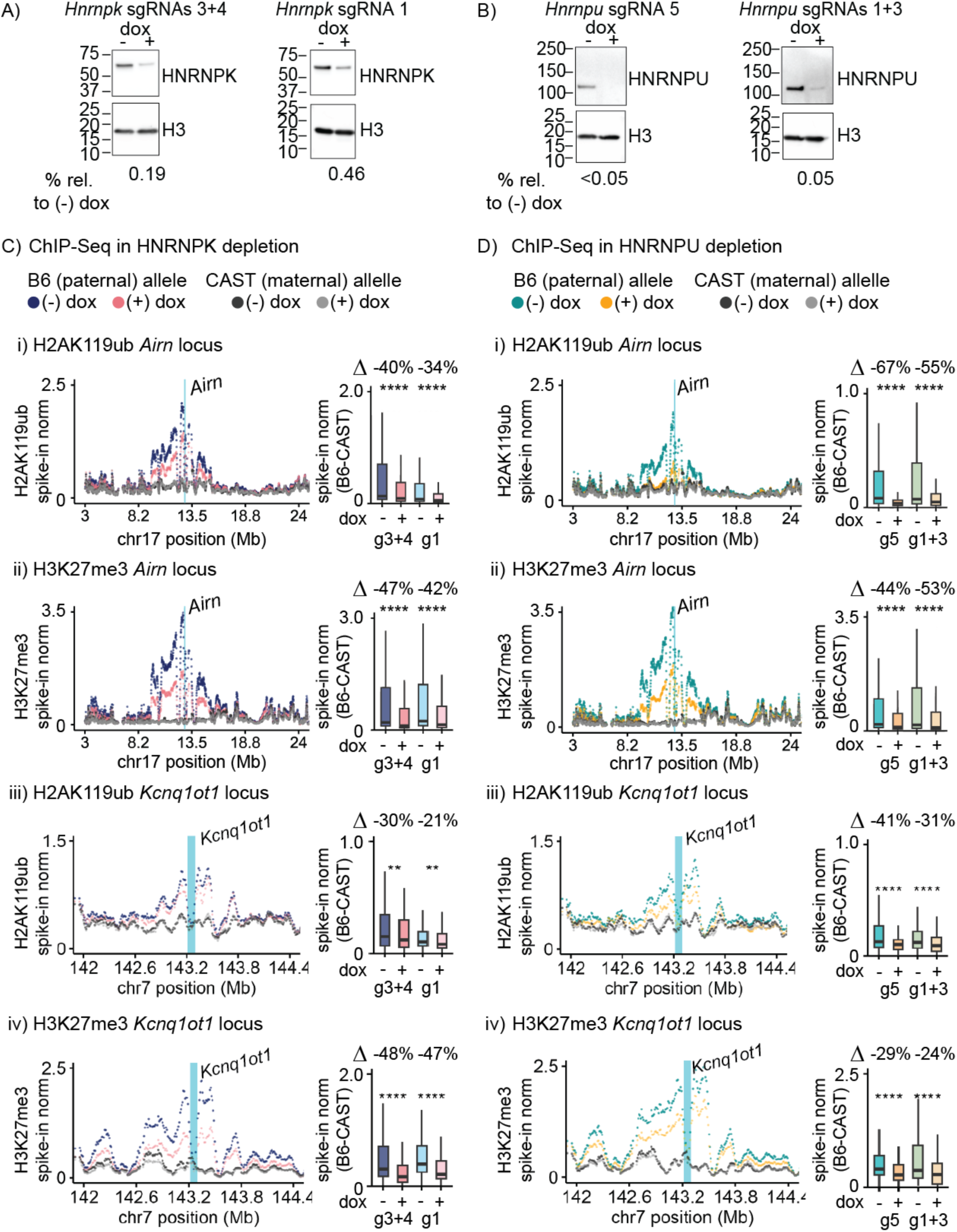
HNRNPU is required for maintenance of PRC-directed modifications induced by *Airn*, *Kcnq1ot1*, and *Xist*. (A, **B)** Western blots of HNRNPK, HNRNPU, H3. **(C, D)** Allelic H2AK119ub and H3K27me3 levels in (-) vs. (+) doxycycline conditions w/sgRNAs targeting HNRNPK (C), HNRNPU (D), over *Airn* (i, ii), *Kcnq1ot1* (iii, iv) domains. Tiling density (left, averaged data) and box-and-whisker plots (right, data by replicate) of spike-in normalized ChIP-seq signal per 10 kb bin. τιvalues, %-fold change of median [B6-CAST] bin between (-) and (+). ****: *p* ≤ 0.0001; Student’s t-test comparing (-) vs. (+). See also Figures S5-8.

### HNRNPU and HNRNPK depletions derepress genes in *Airn* and *Xist* but not *Kcnq1ot1* target domains

To determine whether HNRNPU and HNRNPK are required for gene repression by *Airn*, *Kcnq1ot1*, and *Xist*, we performed RNA-seq in four biological replicates, comparing TSCs before and after doxycycline addition. We then examined extent to which paternal expression bias was altered by HNRNPU and HNRNPK depletion, looking at genes known to be repressed by *Airn* and *Kcnq1ot1*^4^, and at genes that are subject to, weakly, or strongly escape X-chromosome inactivation. Depletion of HNRNPU led to a modest but significant de-repression of *Airn* and *Xist* target genes of all classes but did not affect repression by *Kcnq1ot1* (Figure 5). Depletion of HNRNPK led to a modest de-repression of *Airn* target genes and genes that strongly escape silencing by *Xist* (Figure 5). Therefore, in TSCs, HNRNPU and HNRNPK are required for maintenance of gene repression mediated by *Airn* and *Xist*, but not *Kcnq1ot1*. This trend largely mirrored changes in PRC-directed modifications upon knockdown of HNRNPU and HNRNPK, which were generally larger in the *Airn* and *Xist* than the *Kcnq1ot1* target domains (Figure 5D).

**Figure 5.**
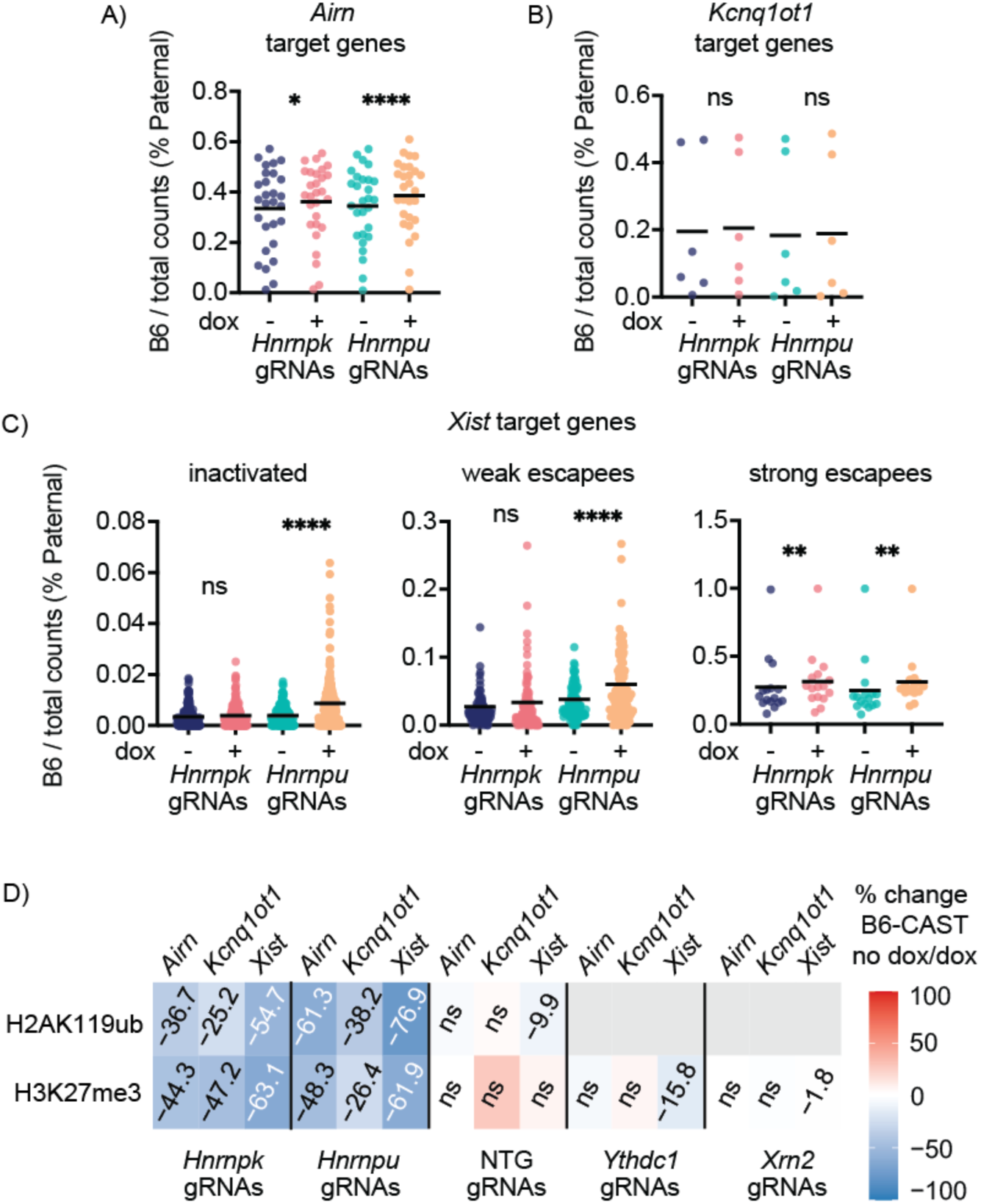
HNRNPU and HNRNPK depletions derepress genes in *Airn* and *Xist* but not *Kcnq1ot1* target domains. (A, B, **C)** %_paternal_expression bias for *Airn* (A), *Kcnq1ot1* (B), *Xist* (C) target genes in (-), (+) doxycycline. Dots, average %_paternal_expression; four independent replicates. *, **, ****: *p* ≤0.05, 0.01, 0.0001, respectively; Student’s t-test. **(D)** Average %-fold change between (-), (+) dox (τιvalues, Figure 4). See also Figure S5-8, Table S4.

### HNRNPK but not HNRNPU depletion reduces associations between PRC1 and *Airn*, *Kcnq1ot1*, and *Xist*

HNRNPK is required to promote accumulation of PRC1- and PRC2-directed modifications within the *Airn*, *Kcnq1ot1*, and *Xist* target domains, presumably by bridging association between PRC1 and regions within each lncRNA.^4,13,34^ We sought to determine whether the requirement for HNRNPU in maintenance of PRC-directed modifications in each lncRNA’s target region could also be explained by reduced associations with PRC1. We performed RIP followed by quantitative PCR (RIP-qPCR) and RIP-seq for RING1B from formaldehyde-crosslinked TSCs after HNRNPK and HNRNPU depletion, as well as in non-depleted controls. By RIP-qPCR, after normalizing for levels of each lncRNA in RNA input, we observed that HNRNPK depletion led to significant reductions in RING1B association with peaked regions in *Airn* and *Xist* (Figure 6A;^13^). In contrast, HNRNPU depletion did not significantly reduce input-normalized levels of association between RING1B and any lncRNA, although we observed a downward trend in *Airn* (Figure 6A). Western blots demonstrated that levels of HNRNPK and RING1B were unaltered by HNRNPU depletion (Figure 6B). HNRNPU depletion did not alter patterns of RING1B association in *Airn*, *Kcnq1ot1*, or *Xist* (Figure 6C). Finally, while patterns of association of HNRNPK and HNRNPU over *Airn* and *Kcnq1ot1* were highly correlated, they were less correlated over *Xist*, where HNRNPK, but not HNRNPU, was enriched in peaks over the same regions that also exhibited peaked association with RING1B (Figure 6C). We conclude that HNRNPU is required for maintenance of PRC-directed modifications induced by *Airn*, *Kcnq1ot1*, and *Xist* via a mechanism at least partly distinct from that of HNRNPK.

**Figure 6.**
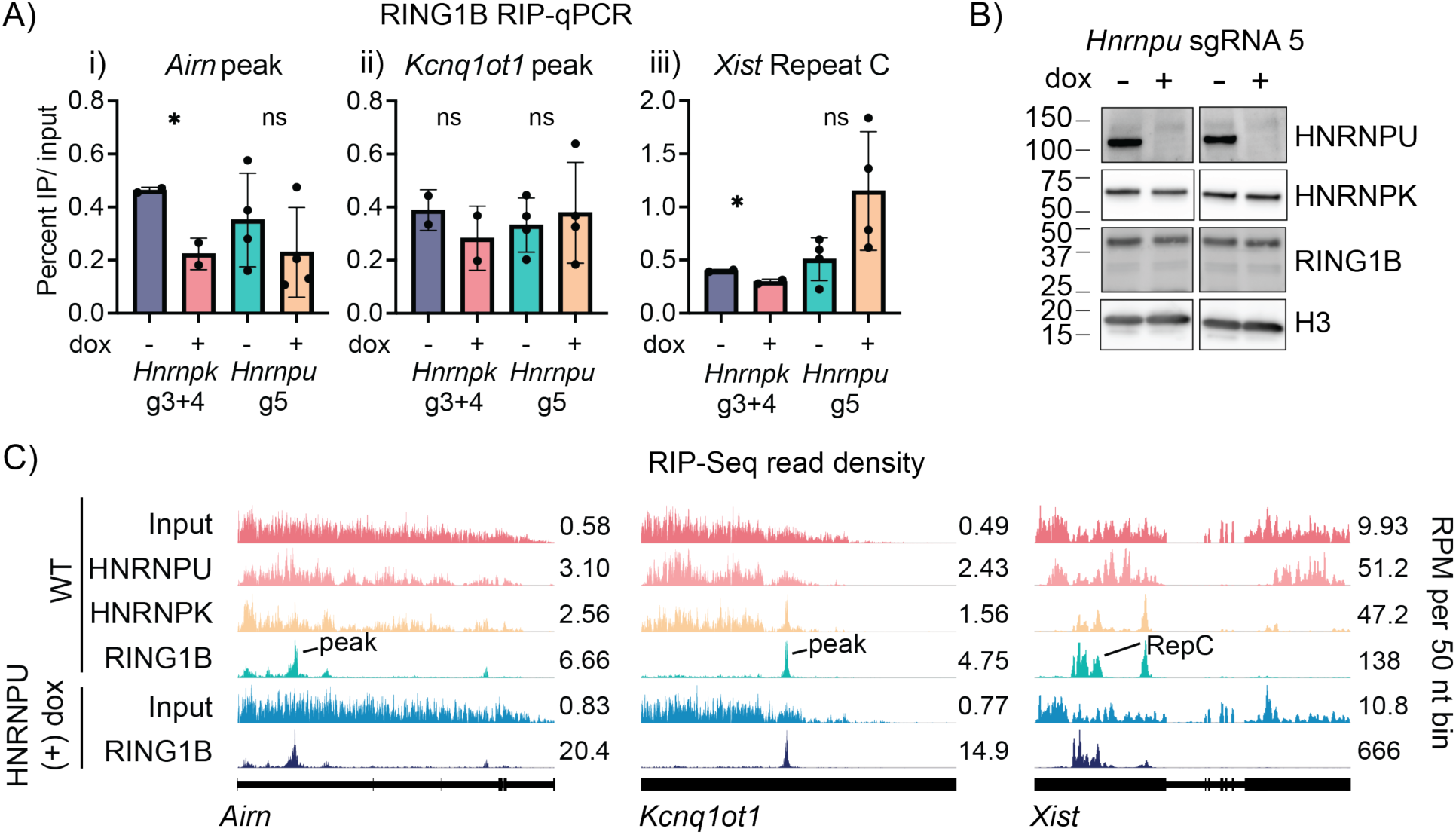
HNRNPK but not HNRNPU depletion reduces associations between PRC1 and *Airn*, *Kcnq1ot1*, and *Xist*. **(A)** Input-normalized RING1B RIP-qPCR in *Airn* (i), *Kcnq1ot1* (ii), *Xist* (iii) peaks in HNRNPK- /HNRNPU-depleted and non-depleted controls. Dots, averaged qPCR technical triplicates from each biological replicate. *, p ≤ 0.05; Student’s t-test. **(B)** Western blots of HNRNPK, HNRNPU, RING1B, H3. **(C)** RIP-seq density profiles of Input, HNRNPU, HNRNPK in WT TSCs (data from Figures 1/2) and HNRNPU-depleted TSCs (data from Figure 4A). “peak/RepC”, regions assessed in (A).

### Divergent effects of HNRNPU depletion on localization and abundance of *Airn*, *Kcnq1ot1*, and *Xist*

HNRNPU promotes association between chromatin and *Xist* and other nuclear-retained RNAs.^62–68^ We therefore sought to determine effects of HNRNPU depletion on localization, abundance, and half-lives of *Airn*, *Kcnq1ot1*, and *Xist*. We used single-molecule-sensitivity RNA FISH to examine lncRNA localization in HNRNPU-depleted versus non-depleted controls. Looking first at *Xist*, after HNRNPU depletion, the primary phenotype observed was a reduced ability to detect RNA FISH signal, consistent with observations from human cells.^66,68^ After setting imaging thresholds to visualize the less-intense signal, we did observe dispersal of *Xist* in HNRNPU-depleted cells, consistent with prior observations (Figures 7A, S9;^62,64–67^). In contrast, while overall sizes of *Airn* and *Kcnq1ot1* RNA FISH foci were qualitatively smaller in HNRNPU-depleted cells, we detected similar numbers of foci for both *Airn* and *Kcnq1ot1* in HNRNPU-depleted and non-depleted cells and did not observe diffusion away from their sites of transcription (Figures 7B, S9). RNA-seq data revealed that HNRNPU and HNRNPK depletion significantly reduced the abundance of *Airn* and *Xist* but not *Kcnq1ot1* (Figure 6C). RNA-seq read densities and half-lives of each lncRNA were unaffected by either protein’s depletion (Figures 7D, 7E). Thus, both HNRNPU and HNRNPK are required to maintain wild-type levels of *Airn* and *Xist* without altering their stability and are not required to maintain levels or stability of *Kcnq1ot1*. Moreover, HNRNPU depletion does not lead to obvious dispersal of *Airn* or *Kcnq1ot1* from their sites of transcription. Therefore, although HNRNPU is required for the accumulation of PRC-directed modifications in genomic regions targeted by *Airn*, *Kcnq1ot1*, and *Xist*, there are divergent effects of its depletion on localization and abundance of each lncRNA.

**Figure 7.**
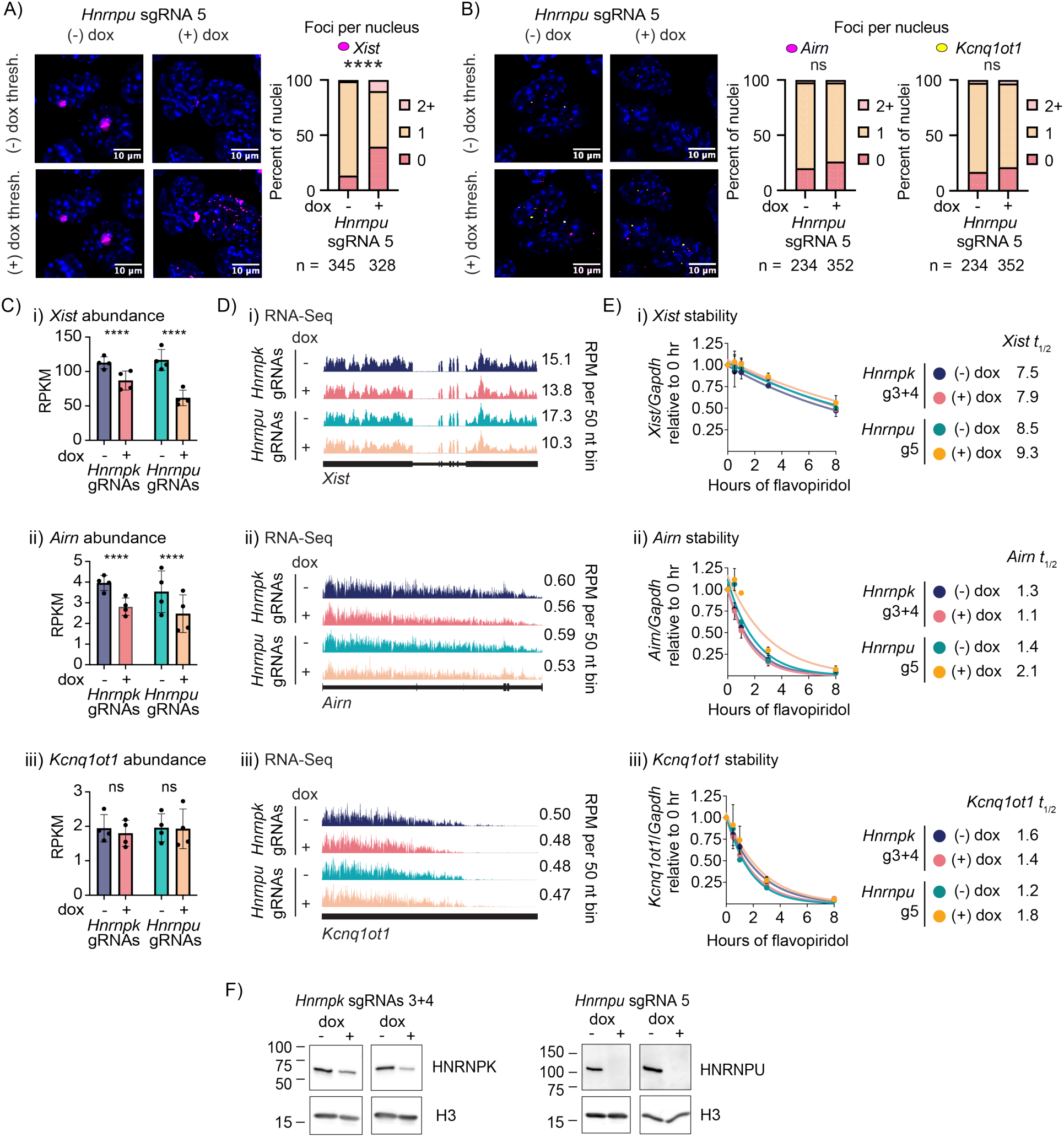
Divergent effects of HNRNPU depletion on localization and abundance of *Airn*, *Kcnq1ot1*, and *Xist*. (A, **B)** RNA FISH detecting (A) *Xist* and (B) *Airn* and *Kcnq1ot1* (magenta and yellow, respectively) in HNRNPU-depleted and non-depleted TSCs. Signal thresholds set against non-depleted TSCs (“(-) dox thresh.”) or HNRNPU-depleted (“(+) dox thresh.”). Bar plots, lncRNA foci per nucleus in (-) and (+) dox. ****, p ≤0.0001; Chi-squared. **(C,D,E)** Expression (C), RNA-seq read density (D), and expression after flavopiridol (E) of *Xist* (i), *Airn* (ii), *Kcnq1ot1* (iii) in (-) dox, (+) dox conditions in TSCs expressing HNRNPK-, HNRNPU-targeting sgRNAs. Dots, individual biological replicates. ****, p ≤0.0001; Student’s t-test. (**F)** Representative western blots. See also Figure S9.

## Discussion

Examining 27 proteins previously shown to associate with *Xist*, we compared enrichments and patterns of protein association across *Airn*, *Kcnq1ot1*, and *Xist*, evolutionarily unrelated lncRNAs that serve as models to understand the regulatory functions of protein-RNA interactions in the nucleus. Despite differing linear sequence, we identified significant enrichments and networked relationships for several proteins across all three lncRNAs and within other chromatin-enriched transcripts, providing perspectives on protein interactions that may coincide with repressive function in RNA. Our work also provides insights into chromatin regulation by HNRNPU, an essential protein whose mutation causes neurodevelopmental disorders.^39–43^

Central to our study was a formaldehyde-based RIP protocol that employs sonication and IP washes borrowed from ChIP.^44^ Relative to native RIP protocols, our protocol differs in its use of formaldehyde, its sonication prior to IP, and its wash conditions.^44,56^ Relative to the formaldehyde-linked RIP protocol upon which ours was originally based (fRIP;^75^), our protocol differs in its manner of crosslinking, its wash conditions, and its addition of ERCC Spike-In controls prior to library preparation.^44,55^ Our typical RIP-seq analyses leverage a pipeline that can be applied downstream of any peak-calling approach, and defines peak regions as those that are enriched above a minimum threshold value in multiple replicates relative to a specified negative control (typically two-fold above IgG, non-specific epitope tag, or protein knockout, in at least two replicates).^44^

Using a standardized set of analysis procedures to compare to published CLIP and CLAP experiments,^35^ we found that our protocol returned higher signal-under-peak values as well as signal-to-noise and post-lysis reassociation ratios that were largely equivalent. Although CLAP outperformed in signal-to-noise, it still recovered non-specific background and post-lysis reassociation (e.g. over *Xist*/*XIST*). In some instances, our version of RIP appeared to outperform CLIP and CLAP (e.g. post-lysis reassociation of HNRNPU RIP peaks vs. human CLIP and CLAP peaks; Figure 1H). While some differences may be epitope-tag related (3xFLAG for RIP versus Halo-V5 for CLIP/CLAP), the overall similarity in outcomes provides evidence that all three approaches can be used to study RNA-protein associations *in vivo*. This conclusion is supported by prior studies which demonstrated that our protocol exhibits robust and knockout-dependent recovery of RNA associated with endogenous protein targets.^52–54^ Likewise, using our same RIP protocol, we arrived at conclusions similar to those made by Guo and colleagues using CLIP and CLAP and by a third study that developed an RNA-editing strategy to identify RNA-protein associations in situ, which collectively observed that direct interactions between RNA and PRC2, if they occur in vivo, are likely to be low frequency events that do not often rise above thresholds of noise.^13,35,76^

Our protocol offers complementary advantages to CLAP in that it does not require epitope-tagging and involves a standard RNA-seq workflow that includes ERCC Spike-In controls, which enable estimates of post-IP RNA recovery and absolute scaling of RIPs relative to non-specific controls or across experimental conditions. Also, because it uses formaldehyde crosslinking, our protocol recovers both direct and protein-bridged interactions, presumably with limited RNA sequence bias, again complementing UV- and editing-based approaches designed to recover direct protein-RNA interactions.^35,76–78^ In studying mechanisms of protein engagement by lncRNAs such as *Xist*, *Airn*, and *Kcnq1ot1*, which presumably enact functions by assembling multilayered ribonucleoprotein complexes, parallel applications of methods that recover direct versus protein-bridged interactions could prove informative.

Examining our data, we found that the spatial patterns of protein associations within *Airn*, *Kcnq1ot1,* and *Xist* were more similar to each other than most other chromatin-enriched transcripts (Figure 3C), supporting the notion that aspects of mechanism or biogenesis are shared among all three lncRNAs. The similarities were particularly striking between *Airn* and *Kcnq1ot1*, which across a 27-antibody panel, exhibited background-corrected protein association patterns that were more similar to each other than to ∼19,000 other chromatin-enriched transcripts. At the same time, there were differences between the lncRNAs. Namely, *Airn* and *Kcnq1ot1* exhibited enrichments for proteins that were not enriched over *Xist* and vice versa. More strikingly, across our 27-antibody panel, modularity of protein association correlated with the genomic range over which *Airn*, *Kcnq1ot1*, and *Xist* recruit PRCs and repress genes. In TSCs, *Xist* exerts repression over ∼165 Mb, *Airn* over ∼15 Mb, and *Kcnq1ot1* over ∼3 Mb.^4,14^ By several metrics, *Xist* partitioned protein associations to a greater degree than *Airn*, which partitioned to a greater degree than *Kcnq1ot1*. While some partitioning in *Xist* may derive from our *Xist*-focused RIPs, we favor a model in which the partitioning directly reflects the local organization of RNA–protein assemblies across each lncRNA. Specifically, a higher degree of partitioning in *Xist* and *Airn* versus *Kcnq1ot1* may be due to the presence of a greater number of domains in the former lncRNAs that coordinate interactions with multiple proteins simultaneously. Indeed, tandem repeat domains in *Xist* are known protein interaction hubs,^25^ and *Airn* and *Kcnq1ot1* are expressed at equivalent levels and have equivalent half-lives in TSCs,^4^ supporting the notion that their differences are driven by RNA sequence and not expression. Collectively, the RNA-protein associations we detected by RIP likely represent a combination of direct and protein-bridged interactions, along with some degree of noise and, in the case of *Xist* and presumably other RNAs harboring ultra-high-affinity RBP binding sites, post-lysis reassociations which may or may not reflect those that occur *in vivo*. Future work is needed to distinguish between these possibilities and determine the mechanisms and significance of protein engagement by *Airn*, *Kcnq1ot1*, and *Xist*.

To that end, based on its enriched association with *Airn*, *Kcnq1ot1*, and *Xist*, its association patterns that correlated with the PRC1 components RING1B and RYBP, and its requirement for *Xist* function,^64–66^ we examined the role of HNRNPU in repression by *Airn* and *Kcnq1ot1*. We observed that HNRNPU was required to maintain long-range PRC-directed modifications induced by all three lncRNAs, and that HNRNPU (and HNRNPK) were required to maintain wild-type levels of gene repression by *Airn* and *Xist*. The reductions in gene repression were weak compared to reductions in PRC-directed modifications, and we speculate that the former arises from the latter. We likewise speculate that the varied effects of HNRNPU/K depletion on gene silencing by *Airn*, *Kcnq1ot1*, and *Xist* may be due to varying effects of PRC-directed or other epigenetic modifications within each lncRNA’s target domain.

Reductions in PRC-directed modifications upon HNRNPU depletion occurred without an apparent loss of association between PRC1 and *Airn*, *Kcnq1ot1*, or *Xist*, suggesting that unlike HNRNPK, HNRNPU does not bridge the lncRNAs with PRC1.^13^ We likewise observed no role for HNRNPU in tethering *Airn* or *Kcnq1ot1* to chromatin, as has been observed for *Xist.*^64–66^ Considered together, our data suggest a revised model for HNRNPU in long-range gene regulation mediated by lncRNAs. HNRNPU plays an architectural role in the nucleus, helping to keep transcribed regions accessible by forming what has been proposed to be a mesh-like network in concert with chromatin-associated RNAs.^62–65,67,79^ Thus, rather than specifically tethering lncRNAs to chromatin, HNRNPU may enable lncRNA-induced, long-range PRC-directed modifications by alternate, non-exclusive mechanisms. On the one hand, via its ability to form mesh-like networks, HNRNPU may keep distal regions of chromatin accessible to components of lncRNA-ribonucleoprotein assemblies that recruit PRCs and repress genes; this same mesh-like network may enable *Xist* to infiltrate chromatin across the inactive X-chromosome, giving the appearance of tethering.^62–65,67,79^ On the other hand, HNRNPU may play an architectural role locally, associating with nascent RNA to maintain appropriate expression of *Airn* and *Xist* in what might otherwise be a naturally repressive environment. Under this model, HNRNPU’s lack of requirement in maintaining levels of *Kcnq1ot1* may be due to the genomic environment in which *Kcnq1ot1* is embedded rather than a differential interaction between HNRNPU and the *Kcnq1ot1* RNA itself; indeed, by expression-normalized ranking, *Kcnq1ot1* and *Airn* associate with HNRNPU to equivalent degrees (Figure 2A). Additional research is needed to investigate these models and their potential relation to HNRNPU-mediated developmental disorders.^39–43^

Lastly, our study provides perspectives on the possible spectrum of chromatin-associated RNAs that might regulate genes or chromatin *in cis*. *Xist* has served as a model to understand mechanisms of RNA-mediated gene silencing since its discovery.^80,81^ However, relative to other chromatin-associated RNAs, the genomic range over which *Xist* functions, its nuclear abundance, and its stability are exceptional (Figure 2A).^4,13^ For those reasons, we might expect other cis-regulatory RNAs to share more features with the somewhat less exceptional *Airn* and *Kcnq1ot1*. In that regard, our observation that protein association profiles of *Airn* and *Kcnq1ot1* most frequently resembled those of nascent transcripts produced from protein-coding genes may be important. Splicing can occur co-transcriptionally and often leads to rapid export of processed RNA.^82^ However, post-transcriptional splicing and delayed export after splicing are common.^82–91^ Thus, in addition to lncRNAs, RNAs produced from protein-coding genes might carry out lncRNA-like regulation prior to clearance from chromatin, and much of this regulation might be mediated by introns, given their aggregate length relative to exons. Indeed, the *TTN* pre-mRNA forms an architectural hub in the nucleus that regulates gene expression at multiple levels, providing precedent for this model.^92^

### Limitations of the study

Many RIPs in this study relied on antibodies raised against endogenous proteins, raising the possibility that off-target antibody binding confounds interpretation of protein-RNA associations. Our sense is that this limitation is most relevant to low-affinity, weakly-enriched target sites, where off-target binding is more likely to rival on-target binding in affinity. Most antibodies from our work have been validated in different ways over many years, including by us above, providing measures of confidence in their specificity. Still, orthogonal experiments, including epitope-tagging or genetic validation, are important for in-depth study of any RNA-protein interaction, especially those hovering near thresholds of signal over noise.^13,35,54^

Another limitation is an inability to know with absolute certainty whether an interaction detected by RIP, CLIP, or CLAP occurs in vivo or purely in extracts, due to post-lysis reassociation. Intriguingly, we observed that across RIP, CLIP, and CLAP peaks, IP- and species-specific signal ratios were correlated, suggesting that post-lysis reassociation can occur owing to high on-target affinity (Figure 1, Table S2). In our view, high-affinity regions are more likely to reflect bona fide in vivo RNA-protein binding events than not, particularly when RNA and cognate RBP are present in the same cellular compartment (e.g. *Xist*, HNRNPK, and HNRNPU), and when IP regimes involve high quality antibodies, robust post-IP washes, and high dynamic range.

## Acknowledgements

This work was supported by NIH grants R01GM136819, R01GM121806, and R35GM153293; NSF grant DBI-2228805; and the Yang Family Biomedical Scholar Fund (J.M.C.). M.M.M. was supported by T32GM119999 and F31HD111292. R.E.C. was supported by T32GM007092 and F31HD103334. E.W.A was supported by F31HD114456 and a Marzluff Graduate Fellowship from the RNA Discovery Center.

## Author Contributions

M.M.M., S.L., and J.M.C. conceived the study; M.M.M., S.L., E.W.A., B.A.P, S.P.B., R.E.C., Z.Z., and J.M.C. performed the experiments and/or analyzed data; M.M.M. and J.M.C. acquired funding; and M.M.M. and J.M.C. wrote the paper with feedback from co-authors.

---

**Figure S1.**
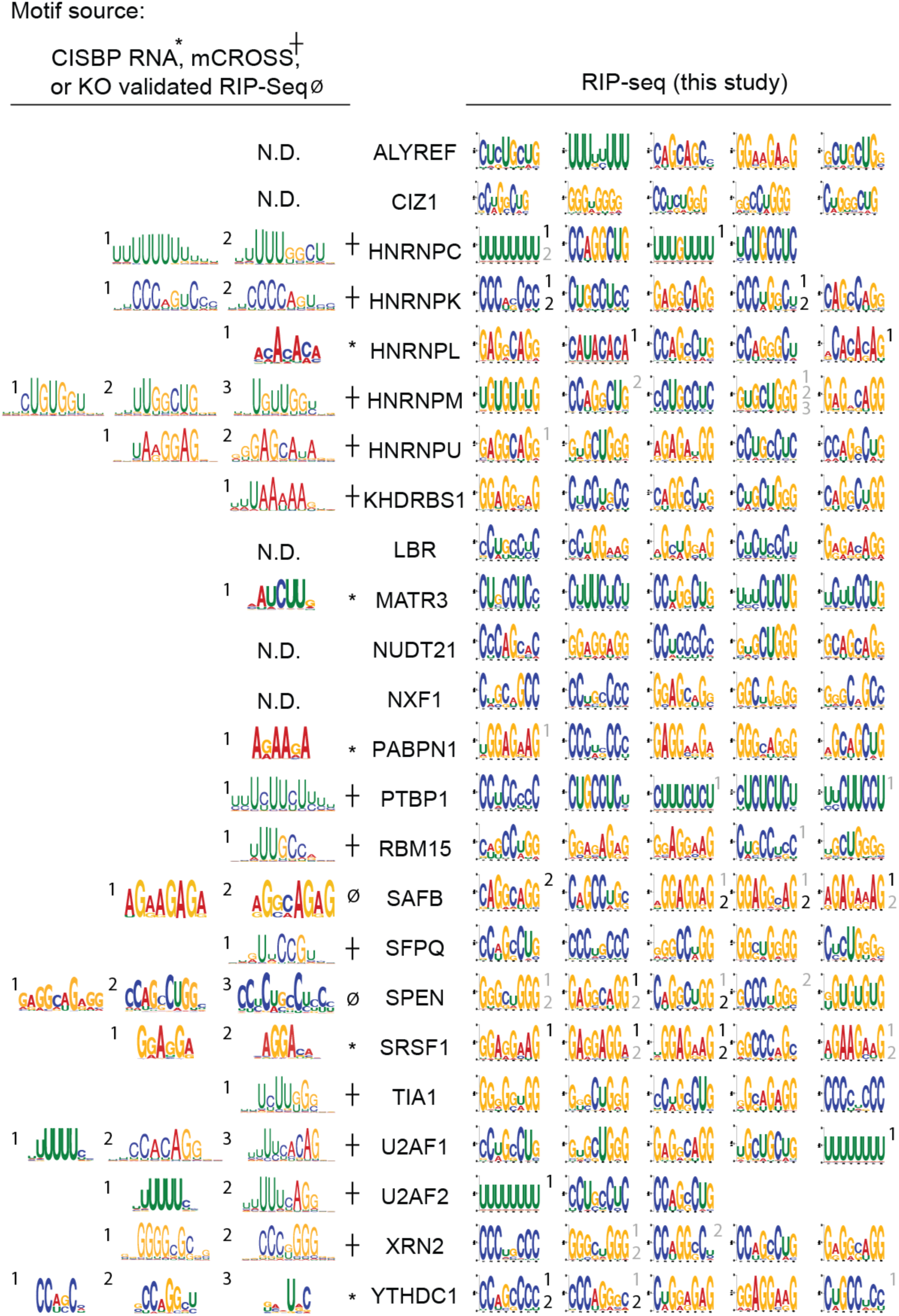
Motifs for RNA-binding proteins derived from RIP-seq in this study versus motifs derived from prior studies (in vitro binding assays, CLIP, or knockout validated RIP-seq). Left-hand panels, motifs reported in CISBP-RNA,^94^ mCROSS,^95^ YTHDC1 CLIP,^73^ or in the case of SAFB and SPEN, motifs derived from studies which used the same antibodies in this study and derived motifs by comparing RIP-seq from wild-type cells versus cells in which the denoted protein was knocked out.^52,53^ Prior CLIP-seq performed to map the RNA targets bound by ALYREF, NXF1, and NUDT21 (a.k.a. CPSF5/CFIm25) failed to identify strong consensus motifs.^96,97^ Right-hand panels, up to the top 5 motifs derived from RIP-seq experiments in this study. Tomtom was used to compare previously identified motifs to those identified in this study ^98^; motifs on the right are marked by the number of the motif on the left with which they share significant similarity. Numbers on the right that are colored in grey signify a p-value of motif similarity of <0.05, and numbers colored in black signify a q-value of <0.05. The p-value of similarity between MATR3 CisBP-RNA motif #1 and RIP motif #5 was 0.057. RING1B, RYBP, and SUPT16H were not included in these analyses.

**Figure S2.**
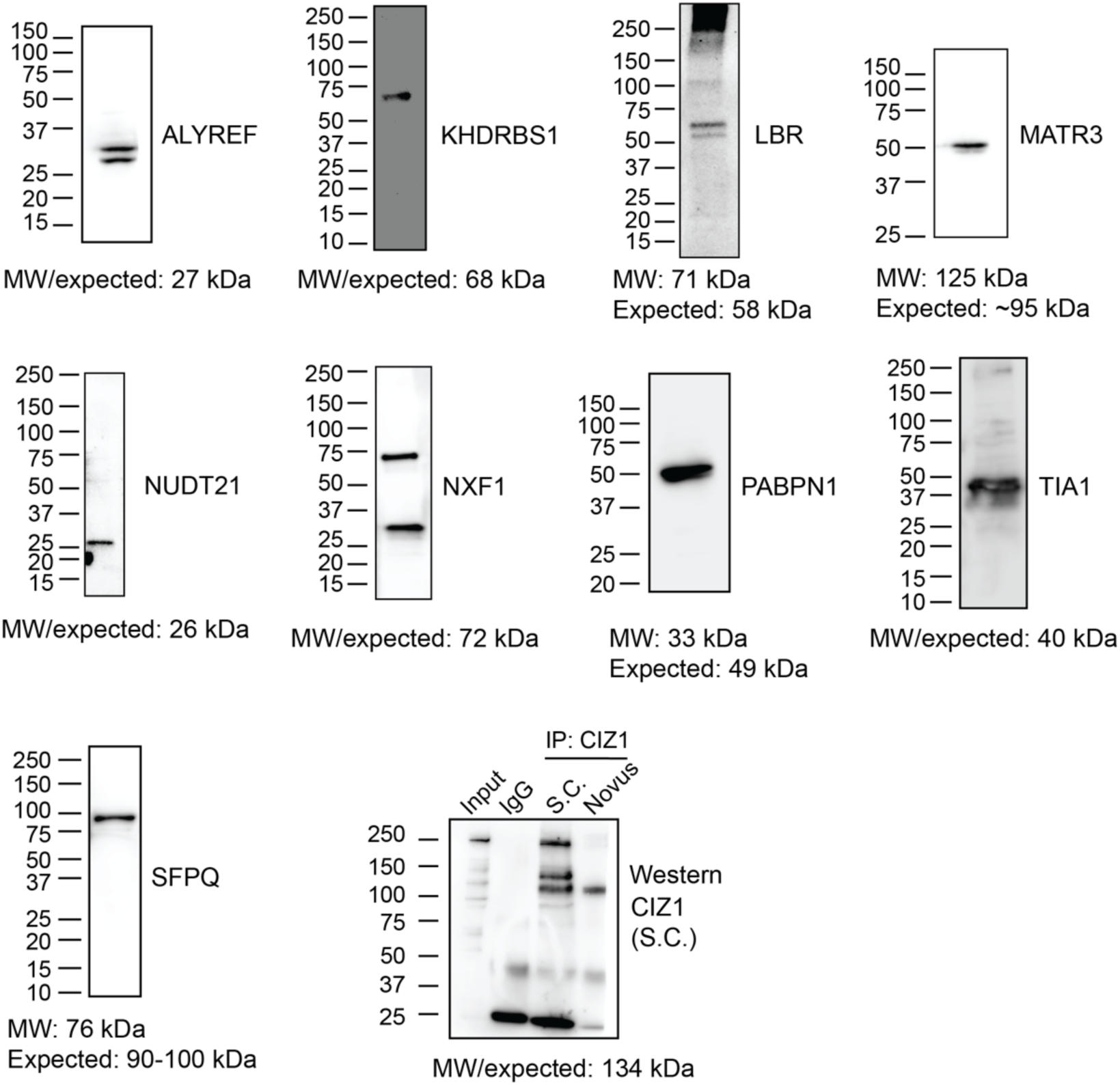
Western blots examining specificity of RIP antibodies raised against RIP-target proteins that lack previously reported consensus RNA interaction motifs or whose RIP-defined motifs lacked significant similarity to CLIP- or *in vitro*-defined motifs. CIZ1 was only detectable by IP-Western; we used the CIZ1 antibody from Novus for RIP-seq in this study. S.C., CIZ1 antibody from Santa Cruz Biotechnology (sc-393021). IP western was performing using RIPseq washes as described.

**Figure S3.**
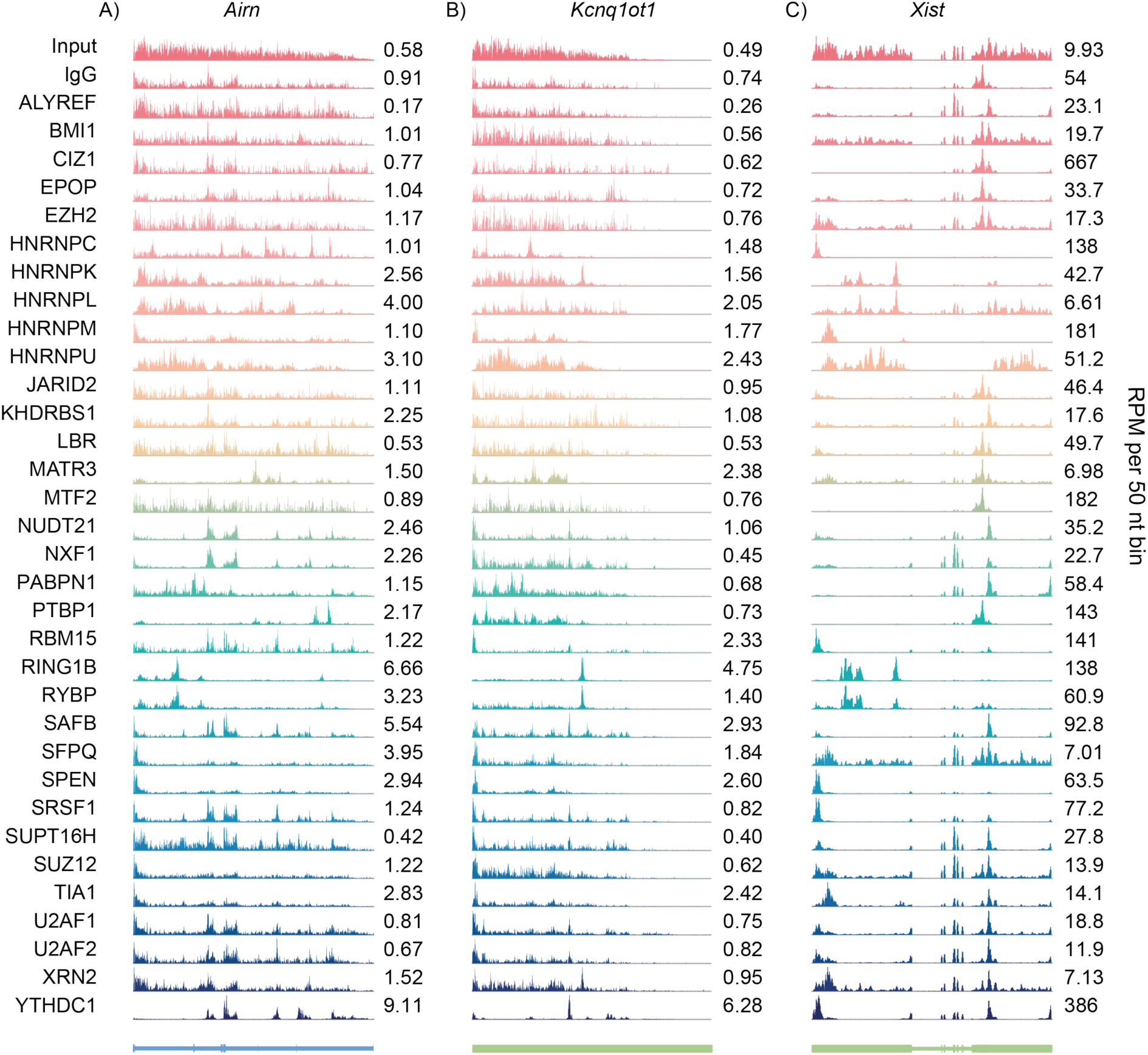
Wiggle density tracks of RIP-seq data over *Airn* (A), *Kcnq1ot1* (B), and *Xist* (C) genes. y-axes, RPM per 50 nt bin.

**Figure S4.**
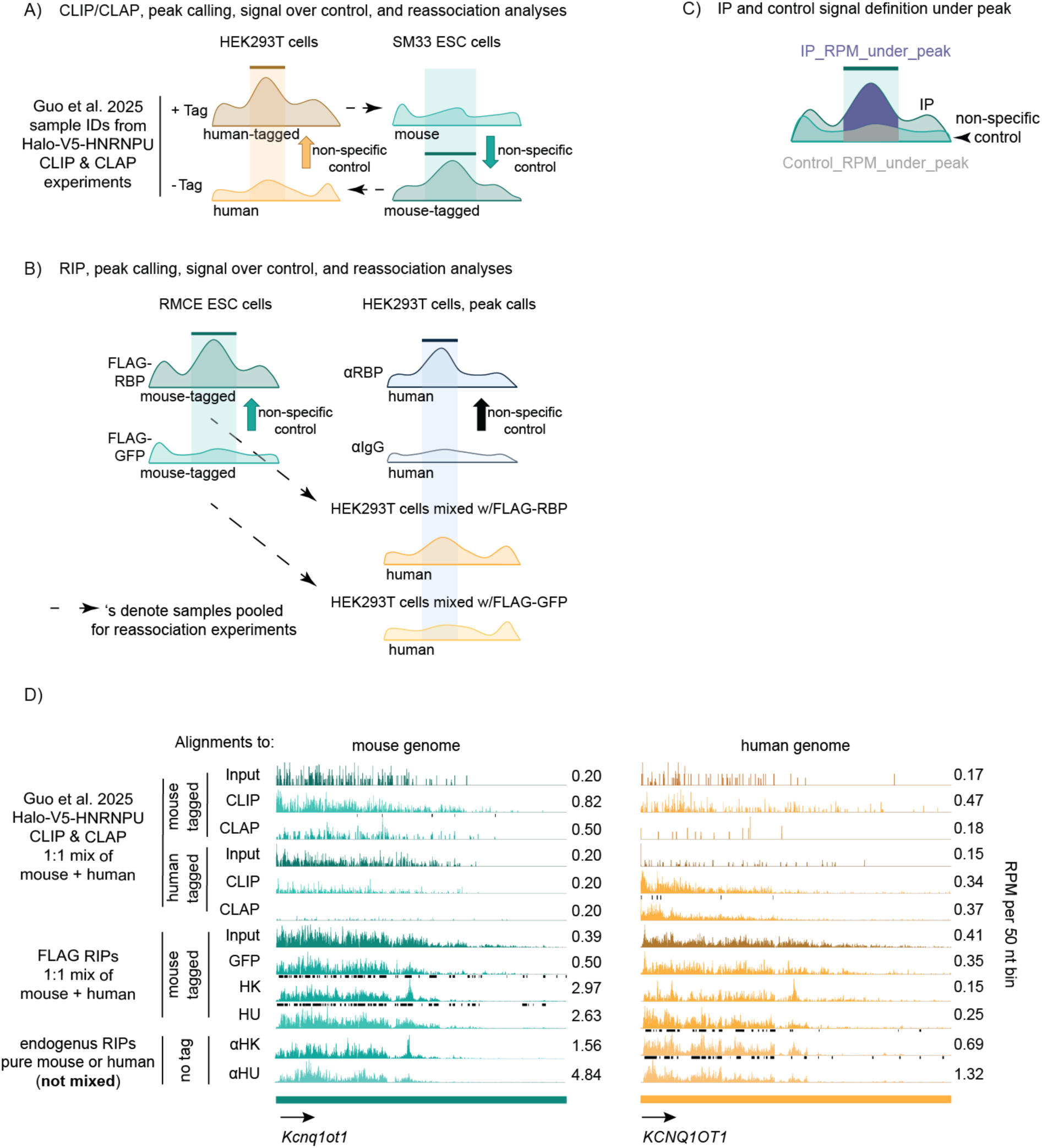
Overview of peak definition and reassociation analyses in RIP, CLIP, and CLAP comparisons. (A,B) Schematics demonstrating peak calling, signal over control, and reassociation strategies for CLIP and CLAP data from Guo and colleagues **(A)**^35^ versus RIP experiments performed in this study **(B)**. In all CLIP and CLAP samples: human 293T and mouse ESCs were mixed in 1:1 ratios. Peaks (shown as thick horizontal bars drawn above enriched regions) were called by comparing signal from the tag-expressing and tag-lacking genomes of the same species. For example, human peaks are called by comparing +Tag samples (which express the tagged RBP in human cells) to -Tag samples (which express the tagged RBP in mouse cells), while mouse peaks are called by comparing -Tag (which express the tagged RBP in mouse cells) samples to +Tag samples (which express the tagged RBP in human cells). Signal was assigned from the tag-expressing genome and noise was assigned from the tag-lacking genome of the same species. Reassociation analyses were conducted by comparing signal under peaks assigned from the tag-expressing genome of the primary species to signal under peaks assigned from the tag-lacking genome of the secondary species. For example, in the mouse *Xist* HNRNPU CLAP reassociation analysis in Figure 1H, CLAP signal under mouse peaks from the mouse-tagged-expressing ESCs (-Tag, third row, lefthand side of Figure 1I) was compared to CLAP signal under human peaks in the tag-lacking human 239T cells that were mixed with the mouse-tagged-expressing ESCs (-Tag, third row, righthand side of Figure 1I). In FLAG RIP samples: FLAG-tag-expressing mouse ESCs were mixed with tag-lacking human 293T cells in 1:1 ratios. Peaks in mouse were called by comparing signal from the FLAG-RBP and FLAG-GFP samples. Peaks in human were called by comparing signal from RBP RIPs using the antibodies raised against endogenous RBPs versus IgG control. Reassociation analyses were conducted by comparing signal under peaks assigned from the tag-expressing mouse genome to signal under peaks assigned from the tag-lacking human genome. For example, in the mouse *Xist* HNRNPU RIP analysis in Figure 1H, RIP signal under mouse peaks from the mouse-tagged-expressing ESCs (tenth row, lefthand side of Figure 1I) was compared to RIP signal under human peaks in the tag-lacking human 239T cells that were mixed with the mouse-tagged-expressing ESCs (tenth row, righthand side of Figure 1I). **(C)** Schematic representing our approach to calculate IP_RPM_under_peak (signal under peak) and Control_RPM_under_peak. **(D)** Wiggle density profiles of RIP, CLIP, and CLAP data over *Kcnq1ot1/ KCNQ1OT1*. Black bars, peaks called in sample. Same format as *Xist/ XIST* data in Figure 1I.

**Figure S5.**
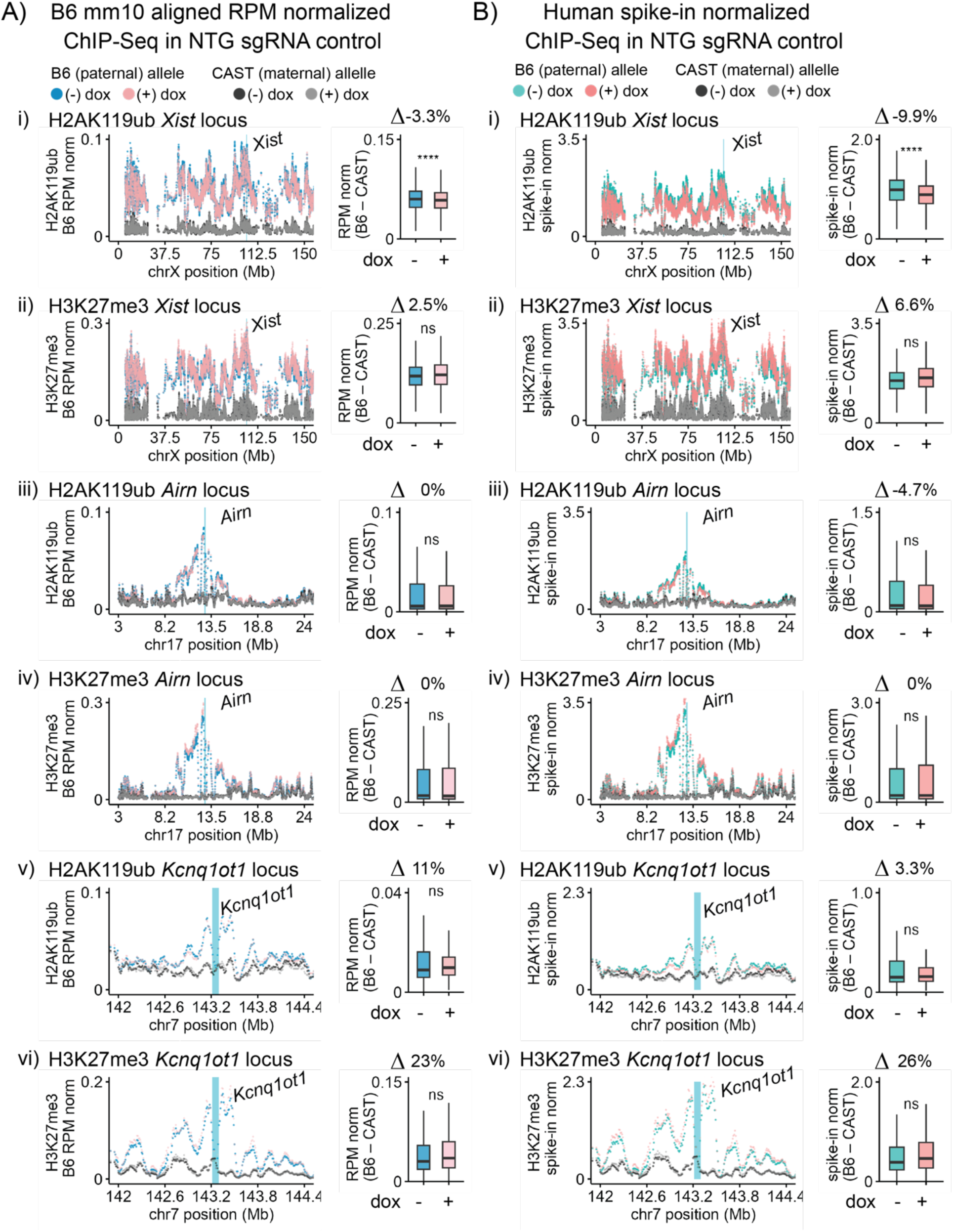
H2AK119ub and H3K27me3 ChIP-seq over *Xist*, *Airn*, and *Kcnq1ot1* target domains in TSCs expressing non-targeting control sgRNA. RPM-normalized (A) and human Spike-In normalized (B) H2AK119ub and H3K27me3 ChIP-seq data from TSCs expressing non-targeting sgRNAs shown over *Xist*, *Airn*, and *Kcnq1ot1* loci. H2AK119ub and H3K27me3 levels are shown in panels (i, iii, v) and (ii, iv, vi), respectively, over the *Xist* (i, ii), *Airn* (iii, iv), and *Kcnq1ot1* (v, vi) target domains, on B6 and CAST alleles with and without Cas9 induction by doxycycline treatment. Tiling density plots and box-and-whisker plots of spike-in normalized H2AK119ub and H3K27me3 ChIP-seq signal per 10 kb bin are displayed on the left and right, respectively, in both (A) and (B). τι values above box plots show the percent fold change of median B6-CAST values between (-) dox and (+) dox. *, **, ****: *p* ≤0.05, 0.01, and 0.0001, respectively; Student’s t-test.

**Figure S6.**
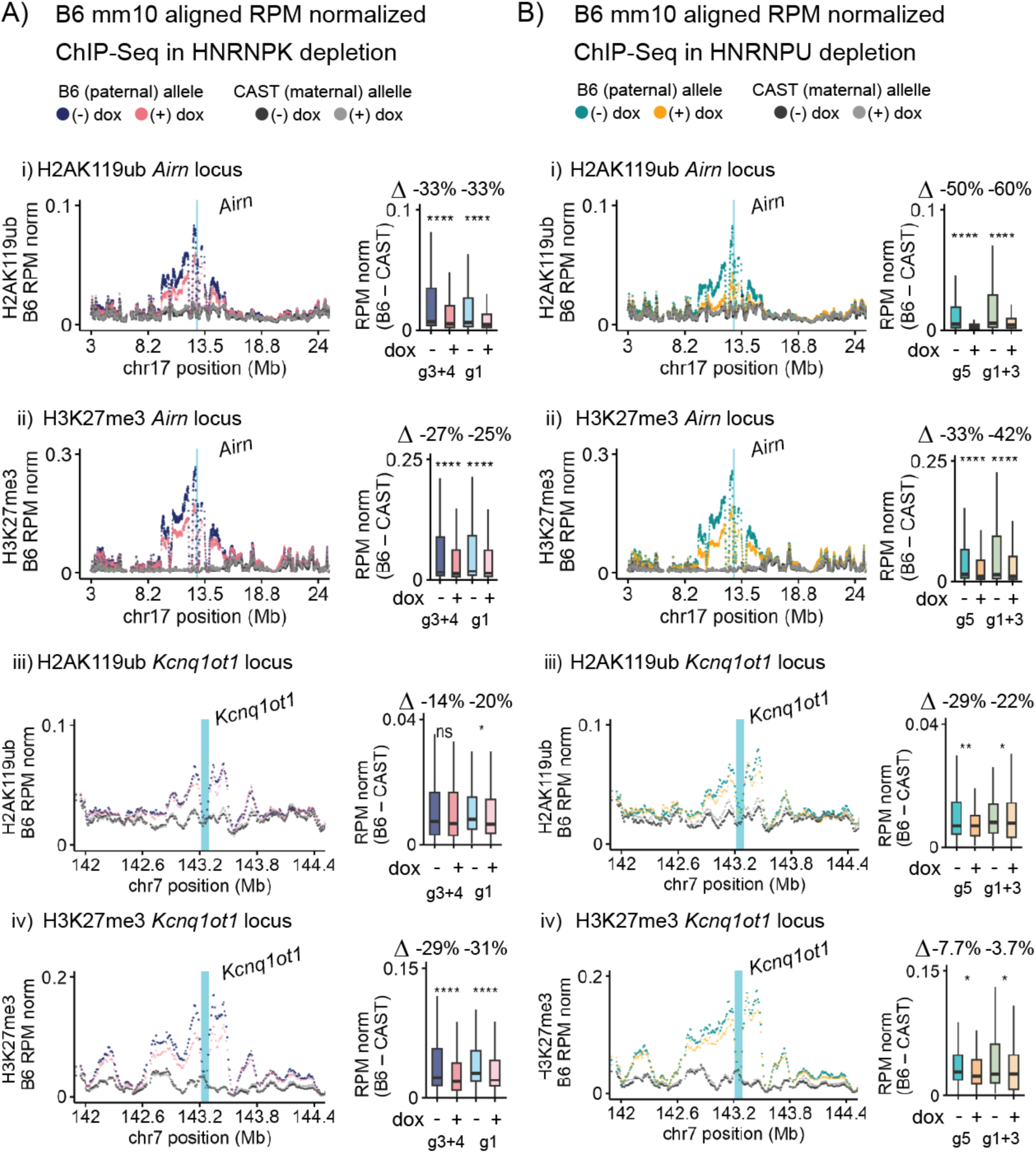
RPM normalized H2AK119ub and H3K27me3 over *Airn* and *Kcnq1ot1* target domains in HNRNPK- and HNRNPU-targeting sgRNA expressing TSCs. RPM-normalized H2AK119ub and H3K27me3 ChIP-seq data in TSCs expressing HNRNPK-targeting (A) and HNRNPU-targeting (B) sgRNAs shown over *Airn* and *Kcnq1ot1* domains. H2AK119ub and H3K27me3 levels are shown in panels (i, iii) and (ii, iv), respectively, over the *Airn* (i, ii) and *Kcnq1ot1* (iii, iv) target domains, on B6 and CAST alleles before and after Cas9 induction with dox. Tiling density plots and box-and-whisker plots of spike-in normalized H2AK119ub and H3K27me3 ChIP-seq signal per 10 kb bin are displayed on the left and right, respectively, in both (A) and (B). Tiling density plots represent the data averaged between the two different sgRNA-expressing populations, and box-and-whisker plots show [B6-CAST] values for each sgRNA experiment. τι values above box plots show the percent fold change of median B6-CAST values between (-) dox and (+) dox for each replicate. *, **, ****: *p* ≤0.05, 0.01, and 0.0001, respectively; Student’s t-test.

**Figure S7.**
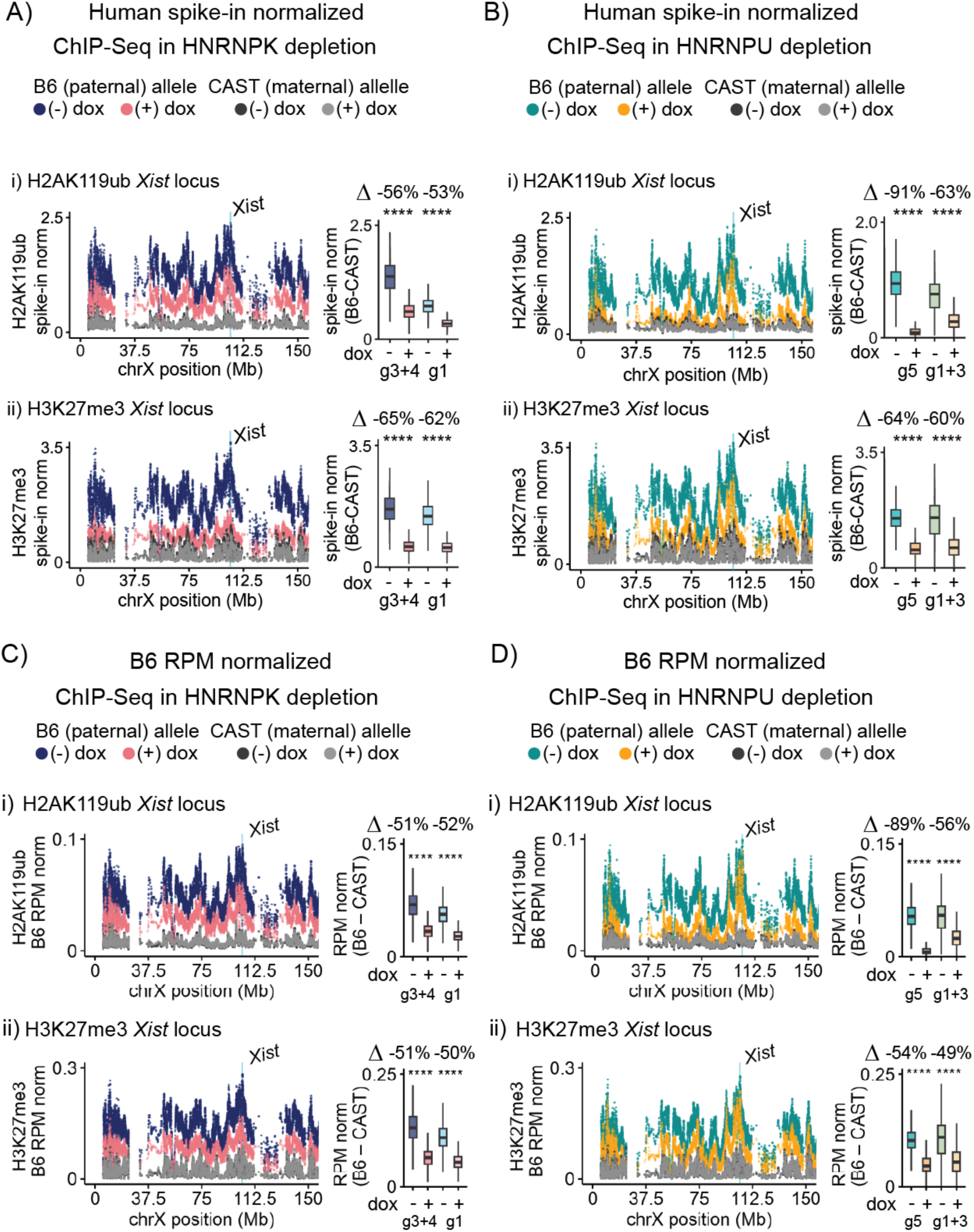
RPM and human spike-in normalized H2AK119ub and H3K27me3 over *Xist* target domain in TSCs expressing HNRNPK- and HNRNPU-targeting sgRNAs. H2AK119ub and H3K27me3 ChIP-seq data in TSCs expressing HNRNPK-targeting (A,C) and HNRNPU-targeting (B,D) sgRNAs shown over *Xist* target domain. Data are shown using both human Spike-In (A,B) and RPM (C,D) normalization schemes. H2AK119ub and H3K27me3 levels are shown in panels (i) and (ii), respectively, on B6 and CAST alleles before and after four days of Cas9 induction with dox. Tiling density plots and box-and-whisker plots of spike-in normalized H2AK119ub and H3K27me3 ChIP-seq signal per 10 kb bin are displayed on the left and right, respectively, in (A-D). Tiling density plots represent the data averaged between the two different sgRNA-expressing populations, and box-and-whisker plots show [B6-CAST] values for each sgRNA experiment. τι values above box plots show the percent fold change of median B6-CAST values between (-) dox and (+) dox for each replicate. *, **, ****: *p* ≤0.05, 0.01, and 0.0001, respectively; Student’s t-test.

**Figure S8.**
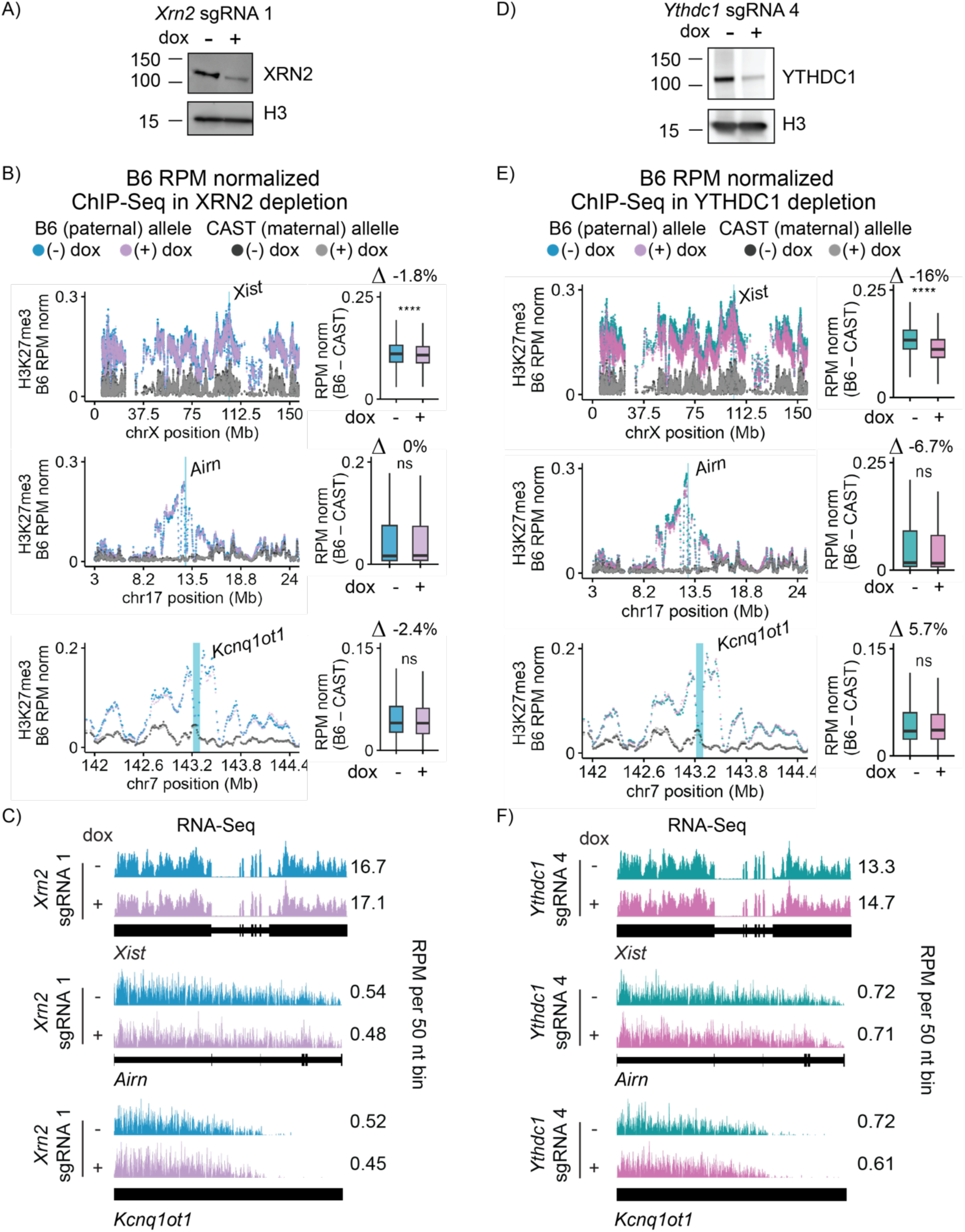
RPM normalized H3K27me3 ChIP-seq over *Xist*, *Airn*, and *Kcnq1ot1* target domains in TSCs expressing XRN2- and YTHDC1-targeting sgRNAs. (**A, D**) Western blot of XRN2 and histone H3 in (-) and (+) dox conditions, in TSCs expressing *Xrn2*-targeting (A) and *Ythdc1*-targeting (D) sgRNAs. **(B, E)** RPM normalized H3K27me3 ChIP-seq data in TSCs expressing *Xrn2*-targeting (B) and *Ythdc1*-targeting (E) sgRNAs, shown over *Xist*, *Airn*, and *Kcnq1ot1* target domains. H3K27me3 levels are shown on B6 and CAST alleles in (-) and (+) dox conditions (four days of Cas9 induction). Tiling density plots and box-and-whisker plots of RPM normalized H3K27me3 ChIP-seq signal per 10 kb bin are displayed on the left and right, respectively, in both (B) and (E). τι values above box plots show the percent fold change of median B6-CAST values between (-) dox and (+) dox. *, **, ****: *p* ≤0.05, 0.01, and 0.0001, respectively; Student’s t-test. Data shown are from a single experiment. **(C, F)** Wiggle density plots showing RNA-seq data over *Xist*, *Airn*, and *Kcnq1ot1* in (-) and (+) dox conditions in TSCs expressing *Xrn2*-targeting (C) and *Ythdc1*-targeting (F) sgRNAs.

**Figure S9.**
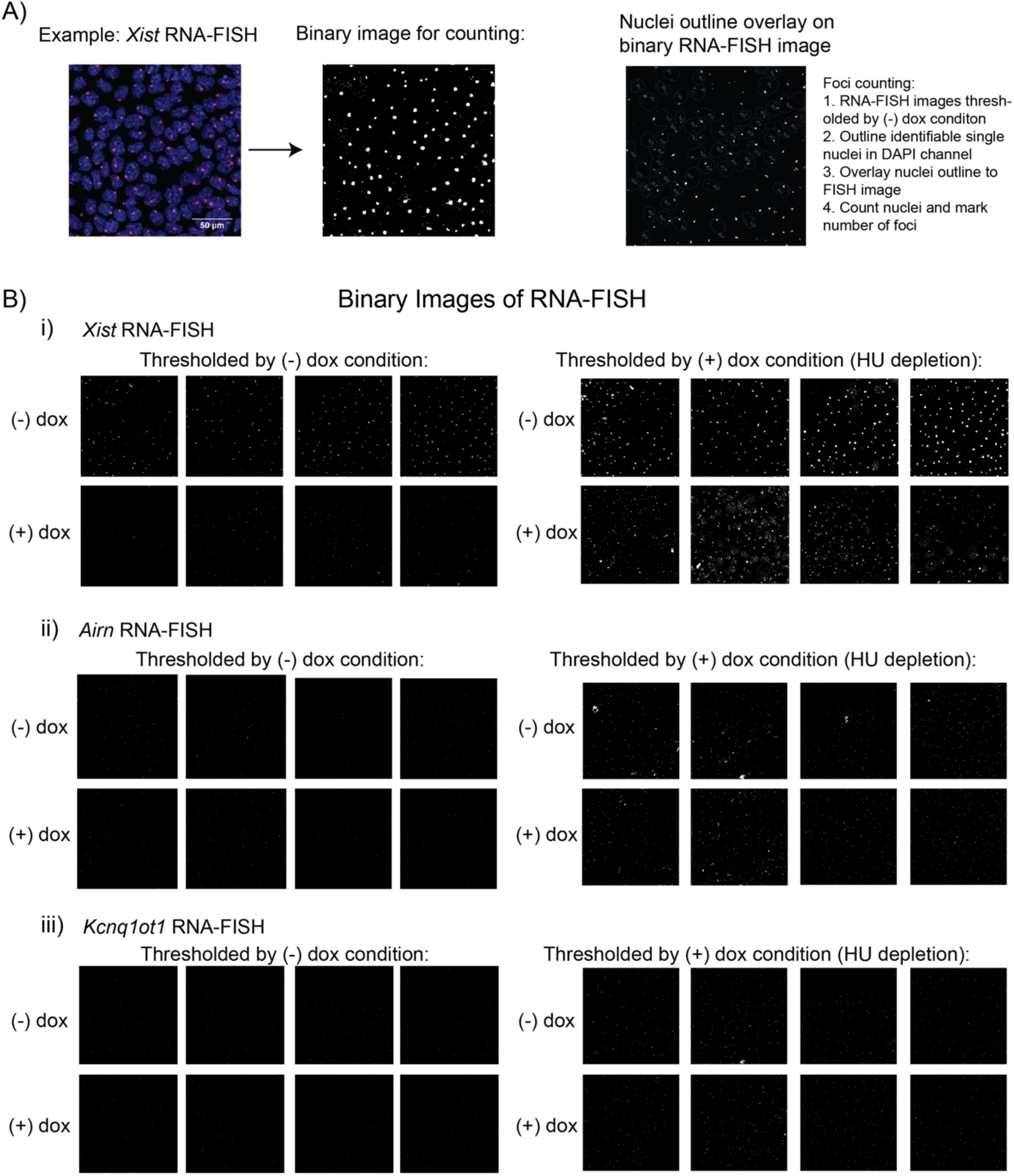
Overview of RNA FISH quantitation strategy and representative images. (A) Overview of foci counting strategy. **(B)** Representative *Xist* (i), *Airn* (ii), and *Kcnq1ot1* (iii) RNA FISH images thresholded by signal in the (-) dox condition (left-hand panels) and (+) dox conditions (right-hand panels).

## Methods

### TSC culture

The C/B TSC line used in this study was derived in.^48^ TSCs were cultured as in.^99^ Briefly, TSCs were cultured on gelatin-coated, pre-plated irradiated mouse embryonic fibroblast (irMEF) feeder cells in TSC media (RPMI [Gibco 11875093], 20% qualified FBS [Gibco 26140079], 0.1 mM penicillin-streptomycin [Gibco 15140122], 1 mM sodium pyruvate [Gibco 11360070], 2 mM L-glutamine [Gibco 25030081], 100 μM β-mercaptoethanol [Sigma-Aldrich 63689]) supplemented with 25 ng/mL FGF4 (Gibco PHG0154) and 1 μg/mL Heparin (Sigma-Aldrich H3149) just before use, at 37°C in a humidified incubator at 5% CO_2_. At passage, TSCs were trypsinized with 0.125% trypsin-EDTA in PBS solution (Gibco 25200-072) for ∼4 min at room temperature and gently dislodged from the plate with a sterile, cotton-plugged Pasteur pipette. To deplete irMEFs from TSCs prior to all harvests, TSCs were pre-plated for 45 min at 37°C on a nongelatinized plate, which was then rinsed twice with a serological pipette, and the solution was transferred to a fresh culture plate. MEF-depleted TSCs were then cultured for three to four days in 70% irMEF-conditioned TSC media supplemented with growth factors prior to harvesting for molecular or genomic assays.

### ESC culture

B6/CAST F1-hybrid *Rosa26*-RMCE ESCs from ^53^ were grown on gelatin-coated plastic dishes in a humidified incubator at 37°C and under 5% CO_2_. Cells were grown in DMEM high glucose plus sodium pyruvate (Gibco 11995-065), 15% ESC-qualified fetal bovine serum (Gibco 26140-079), 0.1mM non-essential amino acids (Gibco 11140-050), 100U/mL penicillin-streptomycin (Gibco 15140-122), 2mM L-glutamine (Gibco 25030-081), 0.1mM β-mercaptoethanol (Sigma-Aldrich 63689), and 1:500 LIF conditioned media produced from Lif-1Cα (COS) cells.

### HEK293T cell culture

HEK293T human embryonic kidney cells were grown in DMEM (Gibco) with 10% Fetal Bovine Serum (Gibco), 1% Pen/Strep (Gibco), and 1% L-glutamine (Gibco). Cells were maintained in incubators set at 37°C and 5% CO_2_. Media was replaced every 3 days.

### Generation of Cas9/sgRNA-expressing TSCs

sgRNAs targeting HNRNPK, HNRNPU, XRN2, and YTHDC1 were designed using the Benchling CRISPR Guide RNA Design tool then cloned into BbsI-digested rtTA-BsmbI piggyBac vector (Addgene plasmid #126028; ^71^) using NEB Quick Ligase (NEB M2200S). HNRNPK sgRNAs are from.^4^ The NTG sgRNA is from.^14^ Plasmids were purified from High-Efficiency 5-alpha competent cells (NEB C2987H) using the PureLink HiPure Plasmid Midiprep Kit (Invitrogen K210004). DNA was quantified via Qubit 2.0 fluorometer (dsDNA broad-range kit, Thermo Fisher Q32853). Cloned sgRNA plasmids are in the process of being deposited in Addgene. Individual or pooled sgRNA vectors were mixed with the TRE-Cas9-Cargo vector (Addgene plasmid #126029; ^71^) and the piggyBac transposase vector from ^100^ at a 1:1:1 ratio in a total of 10 μl. The 10-μL (2.5 ug) plasmid mixtures were mixed with 1 million WT TSCs suspended in 100 μL Neon Buffer R (Invitrogen MPK10025). TSCs were then electroporated using a Neon Transfection System (Invitrogen) with two 30-ms pulse of 950 V (program 5) before seeding onto a well of a 6-well of gamma-irradiated drug-resistant (DR4) irMEF feeder cells (ATCC SCRC-1045) in TSC growth medium lacking penicillin-streptomycin. Media was changed the following morning with 70% conditioned media and growth factors. Starting 24 hours after electroporation, cells were selected with G418 (200 ug/ml; GIBCO) and Hygromycin B (150 ug/ml; GIBCO) for 9 days. TSCs were split every 2-3 days, and media was changed daily with 70% conditioned media, growth factors and antibiotics. Following 9 days of selection, TSCs were then split on to irMEFs with standard TSM and growth factors for expansion.

### Western blots

TSCs were harvested by washing twice with 3 mL ice-cold PBS, scraping into 1 mL ice-cold PBS, and transferring to 1.7-mL tubes. Following centrifugation (1200 × g, 10 min, 4°C), supernatants were removed and cell pellets stored at -80°C. Cell pellets were thawed on ice and lysed by resuspension in 75-150 μL ice-cold RIPA buffer (10 mM Tris-HCl pH 7.5, 140 mM NaCl, 1 mM EDTA, 0.5 mM EGTA, 1% [v/v] IGEPAL CA-630, 0.1% [w/v] sodium deoxycholate, 0.1% [w/v] SDS) supplemented with fresh 1 mM PMSF and 1:100 protease inhibitor cocktail (Sigma P8340-5ML). Cells were thoroughly resuspended using 10 up-down strokes with a P1000 pipette, incubated with rocking for 30 minutes at 4 C, then sonicated on ice using a Sonics Vibracell VCX130 probe-tip sonicator at 30% amplitude (two 10-second pulses with 1-minute rest on ice). Cell debris was removed by centrifugation (16,100 × g, 15 min, 4°C), and cleared lysates were transferred to pre-chilled tubes. Protein concentrations were determined using the Bio-Rad DC Protein Assay (Bio-Rad 5000112) and a standard curve of BSA.

Lysates were diluted 3:1 with 4× Laemmli sample buffer (Bio-Rad 1610747) containing 10% (v/v) β-mercaptoethanol. Equal amounts of protein (20 μg) were loaded onto pre-cast polyacrylamide SDS-PAGE gels (4-20% polyacrylamide, Bio-Rad) selected based on target protein molecular weights and ran in Tris-glycine-SDS running buffer (2.5 mM Tris, 19.2 mM glycine, 0.01% [w/v] SDS, pH 8.3) at 130 V at room temperature. Proteins were transferred to methanol-activated Immobilon-P PVDF membranes (Millipore Sigma IPVH00010), in Tris-glycine-methanol-SDS transfer buffer (2.5 mM Tris, 19.2 mM glycine, 20% [v/v] methanol, 0.001% [w/v] SDS, pH 8.3) at 4°C, either for 1 hour at 100 V or overnight at 20 V. Membranes were blocked for 30 minutes at room temperature with orbital shaking in 1x TBS with 1% Casein blocking buffer (Bio-Rad #1610782). Primary antibodies were applied at concentrations listed in Table S5 and incubated overnight at 4°C with gentle rocking. After washing (one quick TBS-T rinse followed by three 10-minute washes in TBS-T with orbital shaking), membranes were incubated with HRP-conjugated secondary antibodies (1:20,000 dilution, goat anti-mouse HRP (Thermo Fisher A16072); or 1:100,000 dilution, goat anti-rabbit HRP (Thermo Fisher G-21234); or 1:100,000 dilution, rabbit anti-goat HRP (Thermo Fisher 31402)) for 45 minutes at room temperature. Following a final round of TBS-T washes, protein bands were visualized using SuperSignal West Femto ECL substrate (Thermo Fisher 34094) and imaged on a Bio-Rad ChemiDoc system in chemiluminescence mode. Western blots assessing the extent of depletion of protein targets were performed for each (-) and (+) dox replicate in this study.

### RIP-seq

See ^44^ for a line-by-line description of the RIP protocol used in this study, along with many of its standard analytical procedures. Prior to RIP, TSCs were grown to 75-85% confluency for one passage off irMEFs, trypsinized, and counted. TSCs were washed twice in cold 1xPBS and then rotated for 30 min in 10 mL of 0.3% formaldehyde (1 mL 16% methanol-free formaldehyde (Pierce, #28906) in 49 mL 1xPBS) at 4°C. Formaldehyde was quenched with 1.2 mL of 2 M glycine for 5 min at room temperature with rotation. TSCs were washed 3x in cold 1xPBS, then resuspended in 1xPBS at 10 million cells per mL, aliquoted to 10 million cells per 1.7-mL tube and spun down. PBS was aspirated, and the pellets were snap frozen in a dry ice methanol bath and immediately transferred to -80°C. RIPs were performed as in.^13^ Protein A/G agarose beads (25 μL; Santa Cruz sc-2003) were washed three times in blocking buffer (0.5% BSA in 1xPBS). The washed beads were then resuspended in 300 μl of blocking buffer, and 10 µg of antibody was added (see Table S5 for exceptions). The beads and antibodies were then rotated overnight at 4°C. The next day, 10 million cells were resuspended in 500 μL RIPA Buffer (50 mM Tris-HCl, pH 8.0, 1% Triton X-100, 0.5% sodium deoxycholate, 0.1% SDS, 5 mM EDTA, 150 mM KCl) supplemented with 1:100 Protease Inhibitor Cocktail (PIC; Sigma Product #P8340), 2.5 μL SUPERase-In (Thermo Fisher Scientific AM2696) and 0.5 mM DTT and sonicated twice for 30 s on and 1 min off at 30% output using the Sonics Vibracell Sonicator (Model VCX130, Serial# 52223R). Cell lysates were then centrifuged at 15,000 × g for 15 min. Supernatants were transferred to a new tube and diluted 1:2 with 500ul in 500 μL fRIP Buffer (25 mM Tris-HCl pH 7.5, 5 mM EDTA, 0.5% NP-40, 150 mM KCl) supplemented as above with PIC, SUPERase-In, and DTT. The total lysate (25 μL) was saved for input. Antibody-bound beads were washed three times with 1 mL fRIP Buffer. To the washed beads, 500 μl of diluted cell lysate (5 million cell equivalents) was added. The beads and lysate were then rotated overnight at 4°C. The beads were then washed once with 1 mL fRIP Buffer and resuspended in 1 mL PolII ChIP Buffer (50 mM Tris-HCl pH 7.5, 140 mM NaCl, 1 mM EDTA, 1 mM EGTA, 1% Triton X-100, 0.1% sodium deoxycholate, 0.1% sodium dodecyl sulfate) before transferring to a new 1.7-mL tube. Samples were rotated at 4°C for five minutes, spun down at 1200xg, and the supernatant was aspirated. Samples were washed twice more with 1 mL PolII ChIP Buffer, once with 1 mL High Salt ChIP Buffer (50 mM Tris-HCl pH 7.5, 500 mM NaCl, 1 mM EDTA, 1 mM EGTA, 0.1% sodium deoxycholate, 0.1% sodium dodecylsulfate, 1% Triton X-100), and once in 1 mL LiCl Buffer (20 mM Tris pH 8.0, 1 mM EDTA, 250 mM LiCl, 0.5% NP-40, 0.5% sodium deoxycholate); each wash included a five-minute rotation at 4°C. At the final wash, samples were transferred to a new 1.7-mL tube. After the final wash, inputs were thawed on ice and bead samples were resuspended in 100 μL of 1x reverse crosslinking buffer (1x PBS, 2% N-lauroylsarcosine, 10 mM EDTA) supplemented with 1 μL SUPERase-In, 20 μL Proteinase K, and 0.5 mM DTT). Samples were incubated for 1 h at 42°C, 1 h at 55°C, and 30 min at 65°C, and mixed by pipetting every 15 min. To the bead and buffer mixture, 1 mL TRIzol was added, vortexed, and 200 μL of CHCl_3_ was added. Samples were vortexed and spun at 12,000 × *g* for 15 min at 4°C. The aqueous phase was extracted, and one volume of 100% ethanol was added. Samples were vortexed and applied to Zymo-Spin IC Columns (Zymo #R1013) and spun for 30 s at top speed on a benchtop microcentrifuge. Next, 400 μL of RNA Wash Buffer (Zymo #R1013) was added, and the samples were spun at top speed for 30 s. For each sample, 5 μL DNase I and 35 μL of DNA Digestion Buffer (Zymo #R1013) was added directly to the column matrix and incubated at room temperature for 20 min. Next, 400 μL of RNA Prep Buffer (Zymo #R1013) was added, and the columns were spun at top speed for 30 s. Next, 700 μL RNA Wash Buffer (Zymo #R1013) was added, and the columns were spun at top speed for 30 s. Next, 400 μL RNA Wash Buffer was added, and the columns were spun at top speed for 30 s. The flow-through was discarded, and the columns were spun again for 2 min to remove all traces of the wash buffer. Columns were transferred to a clean 1.7-mL tube, 15 μL of ddH2O was added to each column, and after a five-minute incubation, samples were spun at top speed to elute. 9 μL from each RIP-seq replicate was mixed with 1 μL of a 1:250 dilution of ERCC RNA spike-in controls (Thermo Fisher 4456740) and subjected to strand-specific RNA-seq using the KAPA RNA HyperPrep Kit with RiboErase (HMR) (Roche 08098140702) and KAPA Unique Dual-Indexed Adapters (Roche 08861919702) or TruSeq adapters. RNA fragmentation was performed at 85°C for 5 min. Libraries were quantified using a Qubit 2.0 fluorometer (dsDNA broad sensitivity kit, Thermo Fisher Q32850), pooled, and sequenced on Illumina NextSeq500 or Illumina NextSeq1000 platforms.

### Generation and crosslinking of FLAG-cDNA-expressing ESCs

FLAG-HNRNPU, -HNRNPK, and -GFP were all constructed as part of ^53^, and are available on Addgene. One day prior to transfection, ESCs generated in ^53^, which contain a doxycycline-inducible *Xist* in *Rosa26* but have had the adjacent hygromycin B resistance gene removed by Flp recombinase, were seeded at 0.5 × 10^6 cells per well of a six-well plate. The following day, 625 ng of rtTa plasmid, 625 ng of cDNA plasmid, and 1.25 µg of piggyBac transposase from Kirk et al. (2018) (2.5 µg total DNA) were mixed with 5 µL P3000 reagent, 7.5 µL Lipofectamine 3000 reagent, and Opti-MEM media (Gibco #31985-070) to a final volume of 250 µL. The reagents were incubated for 10 min at room temperature before being added to cells with fresh media. After 24 h, cells underwent one week of selection by hygromycin B (50 µg/mL) and G418 (200ug/mL).

### Reassociation RIP-seq experiments

RIPs for the reassociation experiments described in Figure 1 were performed as above, with some exceptions. ESCs containing FLAG- expressing constructs were induced with 1000 ng/mL doxycycline for four days prior to crosslinking. Crosslinking for both ESCs and HEK293Ts was performed as above. 10 μg of FLAG antibody (Sigma F1804) was used for each RIP. A 1:1 volumetric ratio of sonicated extracts made from 5 million human cells (293T) and 5 million mouse cells (ESCs) were combined for a total of 1 mL lysate (in 1:1 RIPA/fRIP buffer, as above). 50 μL of the 1:1 human-mouse lysate was saved for 5% input. Mixed lysate was added to washed, antibody-conjugated beads and RIP-seq was performed as above. HNRNPK and HNRNPU RIPs in 293T cells were performed as described above, using 5 µg of Rabbit IgG (Invitrogen #02-6102) and 5 µL of the same HNRNPK and HNRNPU antibodies used for RIP in TSCs.

### ChIP-seq

ChIP-seq experiments examining effects of protein depletion on chromatin modifications were performed after four days of addition of 1000 ng/mL of doxycycline (Sigma D9891) to sgRNA-expressing cells. (-) and (+) dox TSC cultures were crosslinked on the same day for each replicate performed. Prior to crosslinking for ChIP, TSCs were passaged once off irMEFs. Adhered TSCs were crosslinked with 0.6% formaldehyde (Fisher Scientific, cat #: BP531-500) in RPMI media with 10% FBS for 10 min at room temperature, then quenched with 125 mM glycine for 5 min at room temperature. Crosslinked TSCs were then washed twice with ice-cold PBS and scraped with ice-cold PBS containing 0.05% Tween (Fisher Scientific, cat #: EW-88065-31), PMSF, and PIC (Sigma Aldrich, cat #: P8340). TSCs were then spun at 3,000 × g for 5 min at 4 °C to remove PBS, followed by resuspension in ice-cold PBS with PIC and PMSF and divided into 5-million cell aliquots. All ChIPs were performed using 5 million cells, 5 µL of antibody, and 30 μL of Protein A/G agarose beads (Santa Cruz, cat #: sc-2003). Antibody-conjugated beads were prepared by incubating antibody with beads in 300 μL Blocking Buffer (PBS, 0.5% BSA [Invitrogen, cat #: AM2616]) overnight at 4 °C with rotation. Crosslinked TSCs were resuspended in 1 mL Lysis Buffer 1 (50 mM HEPES pH 7.5, 140 mM NaCl, 1 mM EDTA, 10% glycerol, 0.5% NP-40, 0.25% Triton X-100, PIC) and incubated with rotation for 10 min at 4 °C. Cells were then resuspended in 1 mL Lysis Buffer 2 (10 mM Tris-HCl pH 8.0, 200 mM NaCl, 1 mM EDTA, 0.5 mM EGTA, PIC) for 10 min at room temperature. All buffer removal steps were performed with 5-min 1,200 × g spins at 4 °C. The extracted nuclei pellet was then resuspended and sonicated in 500 μL Lysis Buffer 3 (10 mM Tris-HCl pH 8.0, 100 mM NaCl, 1 mM EDTA, 0.5 mM EGTA, 0.1% sodium-deoxycholate, 0.5% N- lauroylsarcosine, PIC) using a Vibra-Cell VX130 (Sonics) with the following parameters: 8–10 cycles of 30% intensity for 30 s with 1 min of rest on ice between cycles to obtain 100-500 bp fragments. Lysates were then spun for 30 min max speed at 4 °C. The detergent compatible protein assay kit (Biorad 5000113) was used along with BSA protein standards to determine protein quantity of lysates, as well as sonicated chromatin from HEK293T cells crosslinked as above with 0.6% formaldehyde. For each ChIP in each sgRNA genotype, equal amounts of TSC lysate by protein content comparing the (-) and (+) dox conditions was retained. 5% of the standardized TSC lysate amount was then removed, and an equivalent quantity sonicated chromatin from HEK293T cells was added as a spike in control. Lysate volumes were brought to 500ul, Triton X-100 was then added to a final concentration of 1%, and 25 µL was removed to serve as input. Lysates were mixed with pre-conjugated antibody bead mixes and incubated overnight at 4 °C with rotation. The next day, the beads were washed five times with 1 mL RIPA Buffer (50 mM HEPES pH 7.5, 500 mM LiCl, 1 mM EDTA, 1% NP-40, 0.7% sodium deoxycholate, PIC) and once with 1 mL TE, each for 5 min at 4 °C with rotation and spun at 2,000 × g for 2 min for buffer removal. The washed beads were then resuspended in elution buffer (50 mM Tris-HCl pH 8.0, 10 mM EDTA, 1% SDS) and placed on a 65 °C heat block for 17 min with frequent vortexing. ChIP DNA was then reverse crosslinked in 0.5% SDS and 100mM NaCl overnight at 65 °C, followed by a 1 h RNaseA (3 μL; Thermo Scientific, cat #: EN0531) treatment at 37 °C and a 2.5 h Proteinase K (10 μL; Invitrogen, cat #: 25530015) treatment at 56 °C. DNA was then extracted with 1 volume of phenol:chloroform:isoamyl alcohol (Sigma-Aldrich, cat #: P3803) and precipitated with 2 volumes 100% ethanol, 1/10 volume 3M sodium-acetate pH 5.4, and 1/1000 volume linear acrylamide (Invitrogen, cat #: AM9520) overnight at -20 °C. DNA was then precipitated with a 30 min max speed spin at 4 °C, washed once with ice-cold 80% ethanol, and resuspended in TE buffer. ChIP-seq libraries were prepared with NEBNext End Repair Module (NEB, cat #: E6050S), A-tailing by Klenow Fragment (3’/50 exo-; NEB, cat #: M0212S), and TruSeq 6-bp index adaptor ligation by Quick ligase (NEB, cat #: M2200S), and NEBNext High-Fidelity 2X PCR Master Mix (NEB, cat #: M0541S). All DNA size-selection purification steps were performed using AMPure XP beads (Beckman Coulter, cat #: A63880). Single-end, 75- and 100-bp sequencing was performed using Illumina NextSeq 500 and 1000 systems, respectively.

### Total RNA preparation

irMEF feeder-depleted TSCs were washed twice with ice-cold PBS and harvested with 1 mL TRIzol (Thermo Fisher 15596026). Total RNA was prepared via standard TRIzol-chloroform extraction, per manufacturer’s instructions, except for the addition of 4 μL 5 μg/μL linear acrylamide (Thermo Fisher AM9520) to promote precipitation in isopropanol. RNase-free water was added to dried RNA pellets, allowed to incubate overnight at 4°C, and resuspended. RNA concentrations were measured via Qubit 2.0 fluorometer (RNA high sensitivity kit, Thermo Fisher Q32852), and integrity of RNA was confirmed by visualizing rRNA bands on an agarose gel.

### Total RNA-seq

9 μL of 100ng/ul solution of TSC RNA was mixed with 1 μL of a 1:250 dilution of ERCC RNA spike-in controls (Thermo Fisher 4456740), and subject to strand-specific RNA-seq using the KAPA RNA HyperPrep Kit with RiboErase (HMR) (Roche 08098140702) and KAPA Unique Dual-Indexed Adapters (Roche 08861919702) or TruSeq adapters. RNA fragmentation was performed at 94°C for 6 min, diluted adapter stocks were 7 μM, and libraries were amplified using 11 cycles of PCR. Libraries were quantified using a Qubit 2.0 fluorometer (dsDNA broad sensitivity kit, Thermo Fisher Q32850), pooled, and sequenced on Illumina NextSeq500 or Illumina NextSeq1000 platforms.

### Fractionated RNA-seq

The fractionated RNA-seq data analyzed in this study were generated and first reported in ^13^.

### RIP-qPCR

RIP-qPCR was performed as described above for RIP-seq, with some exceptions. Crosslinking was performed as described above for RIP-seq. Protein A/G agarose beads were prepared as described, except that all bead centrifugation steps in this RIP-qPCR protocol were 2000 × g for 1 min; 5 μL of 219 ng/μL anti-RING1B antibody (Cell Signaling 5694S) was used for each RIP. 10-million-cell pellets were thawed on ice, resuspended in 508 μL RIPA Buffer containing PIC, SUPERase-In, and DTT, and sonicated and centrifuged as described. A total of 510 μL of cleared lysate was transferred to new tubes on ice, to which 510 μL fRIP Buffer containing PIC, SUPERase-In, and DTT was added and mixed well. 25 μL of this mixture was removed and stored at -20 °C for later processing as a 5% input sample. Antibody-bound beads were washed twice with 1 mL fRIP Buffer before adding 490 μL lysate mixture (∼5 million cell equivalents) and 490 μL of a 1:1 mixture of RIPA Buffer and fRIP Buffer containing PIC, SUPERase-In, and DTT. Overnight bead-lysate incubation and wash steps were performed as previously described. The input-sample Reverse Crosslinking Buffer was prepared on ice by mixing, for each sample, 33 μL 3X reverse crosslinking buffer (3x PBS, 6% [w/v] N-lauroylsarcosine, 30 mM EDTA), 41 μL RNase-free water, 5 μL 100 mM DTT, 1 μL 20 U/μL SUPERase-In, and 10 μL 20 mg/mL proteinase K (Thermo Fisher 25530049). Bead-sample Reverse Crosslinking Buffer was prepared on ice by mixing, for each sample, 33 μL 3X reverse crosslinking buffer (3x PBS, 6% [w/v] N-lauroylsarcosine, 30 mM EDTA), 66 μL RNase-free water, 5 μL 100 mM DTT, 1 μL 20 U/μL SUPERase-In, and 10 μL 20 mg/mL proteinase K. The 25-μL 5% input samples were thawed on ice and mixed thoroughly by pipetting with 90 μL input-sample reverse crosslinking buffer. To washed beads, 115 μL bead-sample reverse crosslinking buffer was added forcefully to resuspend the beads, but the mixture was not otherwise mixed. Samples were heated as described to reverse crosslinks; bead-containing samples had their beads resuspended every 30 min by forcefully ejecting ∼100 μL of the supernatant onto the settled beads to avoid losing beads to the inside of pipet tips. After heating, RNA was purified with TRIzol and the RNA Clean & Concentrator-5 kit (Zymo R1013) as described above, during which 580 μL of the TRIzol/chloroform aqueous phase was mixed with 580 μL of 100% ethanol.

Reverse transcription was performed using the Applied Biosystems High-Capacity cDNA Reverse Transcription Kit (Thermo Fisher 4368814) to assemble 20-μL random-primed reverse transcription reactions in 0.2-mL PCR strip-tubes on ice, each containing 5 μL of 5% input or 100% RING1B IP RNA sample and 0.5 μL 40 U/μL RNaseOUT RNase Inhibitor (Thermo Fisher 10777019). The reverse transcription was run with the following thermocycler parameters: 25°C for 10 min, 37°C for 120 min, 85°C for 5 min, hold at 4°C. A six-standard curve of 4-fold serial dilutions was constructed from each 5% (-) dox input sample: 8 μL of the previous standard was mixed with 24 μL of nuclease-free water. For each qPCR primer pair, a qPCR master mix was generated, containing for each 10-μL qPCR reaction: 5 μL iTaq Universal SYBR Green Supermix (Bio-Rad 1725124), 3 μL nuclease-free water, 0.5 μL 10 μM forward primer, and 0.5 μL 10 μM reverse primer (see Table S5 for oligonucleotide sequences). Technical-triplicate qPCR was performed with 9 μL qPCR master mix and 1 μL of standard-curve or 2-fold-diluted reverse-transcription product. A standard curve was run on every plate for each primer pair. Plates were firmly sealed with Microseal ’B’ PCR Plate Sealing Film (Bio-Rad MSB1001), mixed several times by inversion and low intensity vortexing, and briefly pulsed down in a centrifuge. Reactions were run on a Bio-Rad C1000 Touch Thermal Cycler equipped with a CFX96 Real-Time System with the following program: 95°C for 10 min, 40 cycles of (95°C for 15 s, 60°C for 30 s, 72°C for 30 s, plateread). Using Bio-Rad CFX manager, first setting “Cq Determination Mode” to “Regression” and then using each primer pair’s standard curve (Cq vs. starting quantity) to convert Cq values to starting quantities for each technical replicate, the percent IP/input value for each RIP sample qPCR triplicate was calculated. qPCR technical triplicates were averaged for each biological replicate and plotted as a single dot in GraphPad Prism. Significance was determined via one-tailed t-test comparing the average of triplicate-averaged biological replicate values of (-) and (+) dox conditions.

### Single-molecule RNA-FISH

Stellaris RNA FISH probes were designed against *Kcnq1ot1* utilizing the Stellaris RNA FISH Probe Designer (LGC, Biosearch Technologies, Petaluma, CA) available online at www.biosearchtech.com/stellarisdesigner (version 4.2; parameters: masking level 5; max number of 48 probes; oligo length 20 nt; min. spacing length 2 nt; Quasar 670). *Kcnq1ot1* probe sequences are listed in Table S5. Probes to *Airn* were designed in (Trotman et al., 2025). Probes to *Xist* were designed by Stellaris (product # SMF-3011-1, Quasar 570). TSCs were hybridized with the Stellaris RNA FISH probes following the manufacturer’s instructions for adherent cells, available online at www.biosearchtech.com/stellarisprotocols. Briefly, following four days of culture in 70% conditioned media with growth factors on coverslips, and with the addition of 1000 ng/mL of doxycycline (Sigma D9891), or no doxycycline, sgRNA-expressing cells were fixed with 3.7% formaldehyde and permeabilized overnight in 70% ethanol. Cells were then incubated in Wash Buffer A with formamide for five minutes at room temperature, before hybridization with 125 nM of *Airn/Kcnq1ot1* or *Xist* probes in hybridization buffer at 37°C for 4 hours, then washed for 30 minutes at 37°C with Wash Buffer A. Cells were then incubated with Wash Buffer A consisting of 5 ng/mL DAPI for 30 minutes at 37°C. Cells were then washed for 5 minutes at room temperature once with Wash Buffer B, then fixed with ProLong™ Glass Antifade Mountant (Thermo Fisher P36982) for 48 hours at room temperature in the dark.

Two biological replicates per (-) dox and (+) dox condition were imaged. Images were taken with a Leica DMi8 inverted brightfield microscope, using Leica Application Suite X software version 3.7.5.24914 and the 63x/1.6 objective (pixel size .104 uM). Fields of cells were chosen by scanning for areas with flat easily identifiable individual nuclei. Acquisition setting for each channel (DAPI, 570, 670) were optimized to avoid saturated pixels (low florescence intensity, and exposure time with log histogram signal not piling up on the right) using the (-) dox sample and kept consistent for the (+) dox sample of that replicate. For each condition, 3-5 Z-stack images with a Z-slice of size 0.2 μm were acquired for each channel sequentially, starting with the highest wavelength channel and moving to the lowest wavelength channel to avoid photobleaching. Z-stacks were set by identifying the limits of the Z position where cells went out of focus. Images were deconvolved with Huygens Essential version 20.04.0p3 64b (Scientific Volume Imaging, The Netherlands, http://svi.nl) using the standard deconvolution profile under batch express. For representative images, Z-series are displayed as maximum z-projections, and (-) dox and (+) dox images were simultaneously adjusted in FIJI (version 2.16.0/1.54p) to have the same threshold, brightness and contrast.

### RIP-seq alignments and peak calling

Reads from individual RIP-seq replicates were aligned to mm10 using STAR ^101^ and filtered for MAPQ of ≥30 using SAMtools.^102^ Code used for peak definition was the same code used in ^13^ and is outlined on https://github.com/CalabreseLab/Airn_Xist_manuscript/tree/main. For each protein profiled by RIP, after alignment and MAPQ of ≥30 filtering, sequencing data from all replicates were concatenated and split into two files using SAMtools,^102^ corresponding to alignments that mapped to the positive and negative strands of the genome, respectively. Using a custom perl script, the strand information within the positive and negative strand alignment files was randomized so as to better match the criteria of the MACS peak caller, which uses the average distance between positive and negative strand alignments to estimate the fragment length.^103^ Putative peaks were called on strand-randomized positive and negative strand alignment files, respectively, using default MACS parameters and not providing a background file.^103^ Peak bed files were converted to SAF format and reads under each putative peak were counted for each RIP replicate as well as for the concatenated set of MAPQ of ≥30 filtered reads from all IgG replicates using featureCounts.^104^ Counts per peak per dataset were converted into RPM by dividing by the total number of sequenced reads per dataset and multiplying by 1 million. We defined as peaks those regions whose RPM-normalized values in the RIPs were >2x those in IgG control in at least two RIP replicates. In cases in which peaks were defined from a single replicate, we required that RPM-normalized values in the single RIP were >2x than those in IgG control.

### Identification of motifs enriched under RIP-seq peaks

For each protein analyzed by RIP-seq, peaks were ranked by the amount of averaged RPM-normalized RIP signal per replicate subtracted by the signal from RPM-normalized IgG control. The top 1000 peaks most enriched in each RIP-seq were selected for motif analyses. The sequences underneath each peak were extracted from mm10 genome using bedtools getfasta.^105^ For each protein analyzed, a set of 1000 control sequences of lengths identical to those in the corresponding peak file was generated, but whose sequence content was randomly generated in python using the mononucleotide frequency of the mm10 primary assembly. Peak sequences were then searched for motifs relative to control sequences using STREME and the options: --maxw 8 --minw 4 --nmotifs 10 –rna.^106^

### Motif comparisons

The motif comparison tool Tomtom^98^ from the MEME suite was used to compare the query motifs generated by STREME in our RIP datasets (right side) and the motifs generated from prior studies (left side) in Figure S1. To generate the target database to search, we first combined all motifs in the left-hand side of Figure S1 into a single file. The RIP-derived motifs on the right-hand side of the Figure were used as queries. Tomtom was run with the code: tomtom -oc output_folder_name query_file target_database. p-values and q-values are reported in the Figure.

### Wiggle track generation

Wiggle tracks were generated using protocols and code from ^107^. Sequencing reads were aligned to the mm10 genome using STAR (v2.7.10a; ^108^). Replicates were downsampled and merged using SAMtools (v1.18; ^102^). Merged BAM files were filtered for MAPQ ≥ 30 and then split by strand with SAMtools and a custom script. Bam files were converted to BED12 format using Bedtools (v2.29; ^105^). Wiggle tracks were generated using a custom script which included normalizing wiggles to total aligned reads. WIG files were converted to BigWig format and then plotted over respective genome coordinates using PlotGardner in RStudio ^109^.

ERCC control median of ratios analysis

ERCC sequence information (ERCC92.fa and ERCC92.gtf) downloaded from the github page associated with ^110^ were used to generate an index against which all relevant RIP samples were aligned using STAR. featureCounts was used to count the number of reads aligning to each ERCC sequence in each RIP. Medians of ratios were calculated in two groups, the first corresponding to RIPs reported in Figure 1B; and the second corresponding to RIPs reported in Figure 1G. A per-sample size factor for each group was generated based on raw counts using the median-of-ratios method. Specifically, for each ERCC control sequence, we calculated the ratio of its raw counts in each RIP replicate to the geometric mean across all RIP samples in the group. The median of these 92 ERCC ratios for each RIP replicate is the un-normalized size factor. Size factors were additionally normalized by calculating: (the arithmetic means of size factor across all Control replicates) / (size factor of each RIP sample). The normalized values are shown as points in Figures 1B and 1G.

### Reassociation analyses

Data from mixed human 293T and mouse SM33-ESC Halo-V5-HNRNPU CLIPs and CLAPs were downloaded from NCBI GEO (GSM8021138-44;^35^), and along with data from our own mixed human 293T and mouse ESC FLAG-GFP, -HNRNPU, and -HNRNPK RIPs, were aligned to concatenated mm10 and hg38 genomes using STAR and the --outFilterMultimapNmax 1 option to only retain uniquely aligned reads^108^. Mouse RIP peaks for FLAG-HNRNPK and FLAG-HNRNPU were called as described above, requiring that RPM-normalized signal in both replicates was >2-fold higher than that of the averaged FLAG-GFP control in the same region of the mouse genome (i.e. putative peak). Human RIP peaks for HNRNPK and HNRNPU were called as described above, using IgG as a control and data collected from RIPs performed in pure 293T extracts (not mixed with mouse). Peaks in CLIP and CLAP datasets were also called as described above, using as the negative controls the alignments from the sample lacking the tagged construct but matched to the species expressing the tagged construct^35^. For example, to call CLIP or CLAP peaks in 293T cells, the RPM values for human alignments using the “minus-tag” samples were used as negative controls. Conversely, to call CLIP or CLAP peaks in SM33-ESCs cells, mouse alignments using the “plus-tag” samples were used as negative controls (the “plus tag” and “minus tag” nomenclature from ^35^ is assigned using a human-centric perspective). See Figure S4.

To rank peaks, we took the product of overall signal strength and the percent of signal represented by IP versus non-specific control, all using the RPM unit value. Specifically, for each peak, we subtracted the averaged RPM of the non-specific control (control_RPM_under_peak) within the peak from the averaged RPM of the IP within the peak, to derive IP_RPM_under_peak, then multiplied that value by the proportion of RPM signal within the peak calculated as [IP_RPM_under_peak]/[RPM_under_peak]. To calculate RIP/CLIP/CLAP signal over *Xist/XIST* and *Kcnq1ot1/KCNQ1OT1*, used the approach described in the “*RIP-seq signal per transcript*” section below.

To calculate species-specific recovery ratios under peaks within each dataset, we calculated RPM values under each peak in the tag-expressing species counting unique alignments to the tag-expressing genome, and separately, calculated RPM values under each peak in the tag-lacking species using alignments to the tag-lacking genome from the same mixed (human plus mouse) alignments described in the first paragraph of this methods section. For each class of peaks analyzed in Figure 1H, we then summed the tag-expressing and tag-lacking RPM values for each peak within the peak class in question, and converted those sums of RPKM values, using the respective peak lengths of each peak class in question. The “species-specific recovery %” values plotted in Figure 1H were derived from the following calculations. In the case of analyzed peaks: a normalized version of ∑RPKM_tag_expressing/(∑RPKM_tag_expressing+ ∑RPKM_tag_lacking). In the case of “All reads in IP”, we used the same equation except we replaced the sums of peak RPKM values with all uniquely aligned reads to the tag_expressing and tag_lacking genomes.

A normalization method (“norm_factor”) is needed in these analyses because mouse-to-human ratios as calculated from RNA Inputs did not equal exactly 1:1 in any RIP, CLIP, or CLAP experiment. Norm_factor was calculated using the RNA-seq data from Input samples, in the following manner:

For each RNA Input sample, we defined **A** as all uniquely mapped reads (or read pairs) in the tag_expressing species and we defined **B** as all uniquely mapped reads (or read pairs) in the tag_lacking species. For each CLIP,CLAP, or RIP sample, we defined **C** as the sum of uniquely aligning reads (in the “all reads in IP” comparisons) or the sum of RPKM values across the specific peak class being analyzed (for example, the top10k ranked peaks) in the tag_expressing species, and we defined **D** as the sum of the analogous peak class in the tag_lacking species. The normalization was then performed as 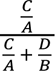. Raw mouse and human read values from all mixing experiments including CLIP and CLAP are reported in Table S1.

### RIP-seq signal per transcript

RIP-seq signal per transcript was calculated as in ^13^. To rank each expressed chromatin-associated transcript by its level of association with each protein profiled by RIP, we first created a version of the GENCODE vM25 basic transcriptome that included one representative intron-containing (i.e., nascent) transcript for each gene that began at the gene’s first annotated transcription start and the last annotated transcription end.^111^ We used kallisto ^112^ with the options “--single -s 300 -l 100 --rf-stranded” to align RNA-seq data from total cellular RNA, as well as cytoplasmic and chromatin-associated fractions. We defined as expressed those transcripts whose median TPM expression value from 8 independent replicates of total RNA-seq was >0.0625 (Table S2). We defined as chromatin-associated any expressed transcript for which [chromatin_TPM]/([chromatin_TPM] +[cytoplasmic_TPM]) was >0.75. We excluded any transcript whose length was less than 500 nt. Total and fractionated data from TSCs are from ^13^; total and fractionated data from ESCs are from ^13,113^.

To calculate RIP signal over each expressed transcript and generate the ranking table shown in Figure 2A, we selected only the subset of reads from each RIP-seq and IgG dataset that aligned under the peaks defined for each protein target of RIP. To do this for a given protein target, RIP-seq and IgG reads were aligned to mm10 using STAR and filtered for MAPQ ≥30 using SAMtools.^101,102^ Samtools view was used to split alignments by strand. For each stranded alignment file, we used samtools view to select the subset of RIP-seq and IgG reads that aligned under each peak. Still using samtools view, we converted bam alignments back into fastq format. Fastq data were aligned with kallisto to the same GENCODE vM25 basic transcriptome used above, which also contained one representative nascent transcript for each gene, using the options [-l 200 -s 50 --rf-stranded].^112^ For each transcript, TPM counts from the IgG datasets were subtracted from TPM counts in RIP-seq datasets, and these IgG-corrected values were used to generate transcript rankings shown in Figure 2. The analogous approach was used to calculate signal and non-specific control data in Figure 1, using the corresponding RIP/CLIP/CLAP datasets and their non-specific controls.

### RIP-seq network analyses

Bedtools multicov was used to calculate replicate-averaged, RPM-normalized read densities in 25 nt bins across the set of 19295 chromatin-associated transcripts in TSCs for each RIP-seq protein target along with IgG controls.^105^ Signal in IgG was subtracted from signal in RIP; negative values were set to 0. Pairwise Pearson’s correlation values for all 27 proteins were calculated between all transcript rows in the resulting matrix. Pearson’s correlations were assigned the value “NA” if they involved signal values of “zero” across all bins in a given transcript. For transcripts that had non-zero signal values in at least one RIP, NA values were replaced with “0” values. Seventy-two transcripts had NA values for all positions across all RIPs and were removed from subsequent network comparisons. To assign RIP protein targets to communities within each transcript’s association network, we required that all RIPs had non-zero values in at least one 25 nt bin; this requirement removed 5464 transcripts and enabled our subsequent calculation of *p* values ascribing likelihood of the prevalence and rarity of edges between protein pairs across the chromatin-associated TSC transcriptome. Communities were assigned using the Leiden algorithm, using only non-negative edge weights/Pearson’s r values in the adjacency matrix. To make the plot in Figure 3Di, we calculated the distribution of edge weights (Pearson’s r values) between all nodes within communities and across different communities. We calculated the modularity of each transcript’s network using the function modularity from the R package igraph ^114^ and calculated silhouette width using the function silhouette in the R package cluster.^115^

To determine whether protein pairs present within the same communities represented prevalent or rare events among the chromatin-associated TSC transcriptome we analyzed networks from the 13831 chromatin-associated transcripts that had at least one non-zero bin value in each RIP. Within these networks, the 27 proteins profiled by RIP-seq could be present in 351 possible pairs (27*26/2). If the network structure is fixed, and node IDs are shuffled to simulate randomization, for each network, the probability that any two proteins fall in the same community (the intracommunity probability) can be estimated as the sum of intracommunity edges divided by the total number of edges in that specific network. Because each network has a different intracommunity probability, (defined below as “prob”), we used an application of the Poisson Binomial Distribution and the cumulative distribution function in the poibin R package to determine if an intracommunity protein pairs that appears x number of times among the 13831 networks were rare [ppoibin (x, prob)] or prevalent [1 - ppoibin(x - 1, prob)].^116^ Due to numerical instability / rounding errors in computing the Poisson binomial cumulative distribution function, *p* values corresponding to values of less than 1.0e-12 were set to < 1.0e-12, before correcting for multiple testing using the Benjamini-Hochberg method.

### Calibrated Allelic ChIP-seq analysis

ChIP-seq and Input reads were aligned to a custom genome consisting of concatenated mm10 and hg38 genomes, and independently, also aligned to a concatenated genomes consisting of a version of mm10 that had been modified to incorporate CAST single-nucleotide polymorphisms (SNPs) downloaded from the Sanger Mouse Genomes Project on 7/30/2020 ^117^ as well as the hg38 genome. Alignments were performed using Bowtie2 with default parameters^118^. Aligned reads that had a MAPQ greater than or equal to 30 were extracted with SAMtools ^102^.

Reads that aligned to uniquely to either the human or mouse were counted. Then mouse reads were retained and human reads were removed from both B6 and CAST alignments. Allele-specific read retention (i.e., reads that overlap at least one B6 or CAST SNP) was performed as in ^48,119^ using a custom Perl script (intersect_reads_snps18.pl). For tiling density plots, B6- or CAST-specific reads were summed in 10 kb using a custom perl script (ase_analyzer8_hDbed.pl). To normalize to human spike-in reads, binned counts of the ChIP were then divided by a “ori ratio” as in ^120^. Ori ratios were calculated in two steps: first, multiplying the spike-in read ratio (human/mouse) from input samples by the target read ratio (mouse/human) from IP samples; second, normalizing to the smallest value between (-) dox and (+) dox conditions. Alternatively, binned counts were divided by the total number of mm10 aligned reads in the dataset and multiplied by a million (i.e., RPM). Normalized counts were then divided by the number of B6/CAST SNPs detected in the bin genomic coordinates (i.e., SNP-norm RPM). Finally, bins were averaged every 9 bins in 1bin increments. For tiling plots with multiple replicates, bins were then averaged among samples. For box plots showing difference in B6 and CAST alleles, bins were averaged every 9 bins in 1bin increments for each allele and CAST reads were subtracted from B6 reads in each bin. The B6-CAST value for bins within the lncRNA target regions identified in ^71^ were then plotted as box plots for individual replicates. Statistical significance of differences for each replicate between the (-) and (+) dox conditions was determined using one-sided Student’s t-tests. All plots were generated using ggplot2 ^121^ in RStudio.

### Allelic RNA-seq analysis

Allelic RNA-seq analysis was performed essentially as described ^13^. RNA-seq reads were aligned to mm10, and independently, aligned to a version of mm10 ^122^ that had been modified to incorporate CAST single-nucleotide polymorphisms (SNPs) downloaded from the Sanger Mouse Genomes Project on 7/30/2020 ^117^. Alignment was performed with STAR (v2.7.9a; ^101^), using multi-sample two-pass mapping and the option “--outFilterMultimapNmax 1” to consider only uniquely mapping reads. Using a custom Perl script (intersect_reads_snps16.pl) aligned reads were parsed to identify reads clearly originating from either the B6 or CAST allele (i.e. reads that overlap at least one B6 or CAST SNP). Reads marked as either B6 or CAST were then assigned to genes using a custom Perl script (ase_analyzer10.pl) and the GTF file gencode.vM25.basic.annotation.gtf,^93^ The ratio of B6/(B6+CAST) reads was calculated for each gene from each sample. The *Airn*- and *Kcnq1ot1*-target genes analyzed in this study were reported in Figure 1 of ^4^. Lists of different X-linked genes analyzed in this study (inactivated, weak escapees, strong escapees) were defined as follows: we required that each gene was represented by an average of at least 5 (B6+CAST) reads in each HNRNPK and HNRNPU (-) and (+) dox replicate (sixteen total replicates). From this list, B6/(B6+CAST) ratio were calculated for genes in the (-) dox replicates from HNRNPK and HNRNPU and average for each gene. Genes whose average, B6/(B6+CAST) ratios were < 0.01 were defined as “inactivated”, those whose ratios were ≥ 0.01 but < 0.10 were defined as “weak escapees”, and those whose ratios were ≥ 0.1 were defined as “strong escapees”. Statistical significance of differences for each gene class between the (-) and (+) dox conditions were determined using average, within-genotype B6/(B6+CAST) ratios and two-tailed Student’s t-tests.

### Quantitation of Airn, Kcnq1ot1, and Xist abundance by RNA-seq

For each sample, RNA-seq reads from each replicate were aligned with STAR (v2.7.9a; ^101^) to the mouse GRCm38 (mm10; ^122^) genome with reference to GTF file gencode.vM25.basic.annotation.gtf ^93^, using multi-sample two-pass alignment that incorporated novel splice junctions discovered among all samples in a given experiment. STAR was run with the option “--outFilterMultimapNmax 1” to consider only uniquely mapped reads. For each sample, reads mapping to *Airn*, *Kcnq1ot1*, and *Xist* were counted using featureCounts (Subread v2.0.4; ^104^) and RPM normalized using the total number of reads uniquely mapped by STAR to the mm10 genome.

### RNA-FISH quantification

RNA FISH signal was quantified in FIJI (version 2.16.0/1.54p) as follows. First, channels were split and maximum z-projections were generated. Then, a threshold was set on the (– dox) projection and applied identically to the paired (+ dox) image to create binary masks of RNA FISH foci (presence/absence as shown in Fig. S8A). For each image, with only the DAPI channel visible, identifiable individual nuclei were manually outlined using the FIJI selection tool. These outlines were then saved as regions of interest (ROIs) as an overlay (example shown in Figure S9A, right). Then, the overlay was applied to the corresponding binary RNA FISH image and as each nucleus was counted and labeled, the number of discrete foci (0, 1, or ≥ 2) for each nucleus was recorded. At least 3 fields per condition ((-) and (+) dox) were analyzed among two biological replicates, fixed and imaged on separate days, for a total of > 200 nuclei per condition. The fraction of nuclei in each foci category (0, 1, ≥ 2) was plotted in GraphPad Prism and statistical significance assessed by chi square test.

### Half-life Analysis

RNA half-lives were measured using a timecourse of flavopiridol (Alvocidib) (Selleck chemicals S1230). MEF-depleted Cas9/sgRNA-expressing TSCs were induced for 3 days with 1000 ng/mL doxycycline prior to start of the flavopiridol time course, for a total of 4 days. TSCs were then treated with 1 uM flavopiridol (24 μl of 50 ug/ml flavopiridol in DMSO added to 3 mL of media) for 30 minutes, 1 hour, 3 hours, 8 hours, (or without flavopiridol, “0 h”) and lysed with TRIzol. To measure levels of *Airn*, *Kcnq1ot1*, *Xist*, and *Gapdh* at each time point, RNA was extracted and subjected to RT-qPCR as above, with the following exceptions: cDNA was diluted 1:20 in nuclease-free water and qPCR was carried out with 4 µL of diluted cDNA and 6 µL of master mix, composed of 5 µL iTaq Universal SYBR Green Supermix (Bio-Rad 1725124) and 1 µL of a 5 µM primer mix per well (see Table S5 for oligonucleotide sequences). RNA levels were normalized to *Gapdh* at each time point and calculated as the percentage of RNA relative to the 0 h time point. For (-) dox HNRNPU depletion cells, only one biological replicate was used, as the 0 h sample was compromised. To estimate half-life, *Gapdh*-normalized data for each replicate were averaged and fit to a non-linear one-phase decay model in GraphPad Prism using the equation Y=(Y0) ∗exp(-K∗X). Error bars represent the standard deviation between biological replicates.

### Data deposition

RIP-, ChIP-, and RNA-seq data from this study have been deposited in GEO, under the accession numbers (and reviewer access tokens): GSE299584(arixcokybzcvnkh), GSE299585 (edyrwiowhbyntub), GSE299587 (gfohgwkwjfghfgj).

Code has been posted in GitHub: https://github.com/CalabreseLab/AKX_HNRNPU_manuscript.

Data in Figures have been deposited into Mendeley.

https://data.mendeley.com/preview/3b3xys2743?a=622e79bc-637f-4c7f-9506-31b022f7a155

